# Metagenomic investigation of the equine faecal microbiome reveals extensive taxonomic and functional diversity

**DOI:** 10.1101/2021.04.30.442084

**Authors:** Rachel Gilroy, Joy Leng, Anuradha Ravi, Evelien M. Adriaenssens, Aharon Oren, David Baker, Roberto M. La Ragione, Christopher Proudman, Mark J. Pallen

**Affiliations:** Quadram Institute Bioscience, Norwich Research Park, Norwich, UK; School of Veterinary Medicine, University of Surrey, Guildford, Surrey, UK; The Institute of Life Sciences, The Hebrew University of Jerusalem, Edmond J. Safra Campus, Jerusalem, Israel; University of East Anglia, Norwich Research Park, Norwich, UK

## Abstract

**Background:** The horse plays crucial roles across the globe, including in horseracing, as a working and companion animal and as a food animal. The horse hindgut microbiome makes a key contribution in turning a high fiber diet into body mass and horsepower. However, despite its importance, the horse hindgut microbiome remains largely undefined. Here, we applied culture-independent shotgun metagenomics to thoroughbred equine faecal samples to deliver novel insights into this complex microbial community.

**Results:** We performed metagenomic sequencing on five equine faecal samples to construct 123 high- or medium-quality metagenome-assembled genomes from Bacteria and Archaea. In addition, we recovered nearly 200 bacteriophage genomes. We document surprising taxonomic and functional diversity, encompassing dozens of novel or unnamed bacterial genera and species, to which we have assigned new *Candidatus* names. Many of these genera are conserved across a range of mammalian gut microbiomes.

**Conclusions:** Our metagenomic analyses provide new insights into the bacterial, archaeal and bacteriophage components of the horse gut microbiome. The resulting datasets provide a key resource for future high-resolution taxonomic and functional studies on the equine gut microbiome.

## Introduction

The horse has played a crucial role in human development and in the spread of human populations (1). Domestication of the horse began at least 6,000 years ago and led to diversification into numerous breeds, accompanied by significant biological changes (2). The horse remains an important component of human society, with around 60 million horses worldwide (3). Horses play roles as working animals across the globe, in transport, agriculture or policing. Horse-riding and equine-assisted therapy provide health benefits, while the horse remains an important food animal globally, with 5 million animals slaughtered for food each year and horsemeat now in favour as a low-methane red-meat alternative to beef (4). In the UK, there are around 374,000 horse-owning households and horseracing is the second most attended sport in the country after football, contributing £4.7 billion to the UK economy (5).

As a foraging herbivore, the horse relies on a cellulose-rich diet of grass and legumes. However, unlike cattle, horses have no rumen to digest complex carbohydrates. Instead, they rely on hindgut fermentation: an efficient but enigmatic process—far less well understood than ruminal digestion—that relies on a rich microbial community, the hindgut microbiome, encompassing bacteria, archaea and viruses, together with fungi and other eukaryotic microbes (6–8). This ecosystem plays key roles in nutrient assimilation and feed conversion—effectively turning grass into horseflesh and horsepower. The horse gut also acts as a reservoir of human and equine pathogens and of antimicrobial resistance (9).

Crucially, a range of diseases are known to be associated with disturbances in hindgut microbial ecology, including foal diarrhoea, colitis, laminitis, colic and equine grass sickness (10). Thus, by better understanding the equine hindgut microbiome, we stand to inform interventions that can improve the health and welfare, performance, value and longevity of horses.

Previous studies of the horse hindgut microbiome have documented a rich variety of microorganisms (spanning phyla from all three domains of life) and have shown that the taxonomic composition of this community varies with age, breed, disease status and has changed during domestication (6, 7, 10–17). However, earlier studies have largely relied on short-read meta-barcoding analyses of 16S rRNA gene sequences, which are limited in that they fail to provide resolution down to the species or strain level, reveal nothing about the population structures or functional repertoires of microbial species and fail to cover viruses and eukaryotes. Thus, despite previous efforts—and drawing on comparisons with the human microbiome, where new species are still being discovered (18, 19) —the horse hindgut microbiome presents us with a vast unexplored landscape of taxonomic, ecological and functional diversity, certain to encompass important, yet undiscovered roles. As in studies of the human gut microbiome, faeces provide ready non-invasive access to the gut contents. As part of the Alborada Well Foal study, a cohort study of foal gut microbial development and health in later life, we applied shotgun metagenomics to five equine faecal samples from 12-month-old thoroughbreds to expand our knowledge of this landscape.

## Materials and methods

### Sample collection and storage

Faecal samples were from five, 12-month-old Thoroughbred racehorses from the same location in Ireland. Samples were collected as part of the Alborada Well Foal study, under the University of Surrey’s ethical review framework, project code: NERA-2017-007-SVM. All horses were at pasture when sampled. 100 g of freshly evacuated faeces was collected from each horse in sterile bijous before immediate storage at 4°C on site at the stud. Once shipped, faecal samples were aliquoted and stored at -80 °C until DNA extraction. Samples were thawed and mixed before DNA extraction using the DNeasy PowerSoil kit (Qiagen), following manufacturer’s instructions. Extracted DNA was stored at -20 °C before further analysis.

### Metagenomic sequencing and processing

Illumina sequencing libraries were constructed as previously described by Ravi and colleagues (2019) (20). Paired-end metagenomic sequencing was performed on the Illumina NextSeq, before bioinformatic processing on the Cloud Infrastructure for Microbial Bioinformatics (CLIMB) (21). Output reads (2x150bp) were assessed for quality using FastQC v0.11.8 and then trimmed using Trimmomatic v0.36 configured to a minimum read length of 40 (22, 23). All metagenomic samples described here can be accessed on the Sequence Read Archive under BioProject ID PRJNA590977. Reads were aligned to the horse genome (GCF_002863925.1) using Bowtie2 v2.3.4.1 (24), allowing removal of host reads with SAMtools v1.3.1 (25).

Taxonomic profiling of sequencing reads was performed using Kraken 2 (26) to search a microbial database built from archaeal, bacterial, fungal, protozoan, viral and univec_core sequences in Refseq in January 2020. Bracken was used to estimate taxon abundance from Kraken 2 profiles, accepting only those taxa with >1000 assigned reads (27). Bracken-database files were generated using “bracken-build” on our microbial database and visualised using Pavian (28).

### Metagenomic assembly and binning

Host-depleted reads were assembled individually from each metagenomic sample with MegaHIT (29), using kmer sizes 25,43,67,87 & 101, before assessing the quality of resulting contiguous sequences (contigs) with anvi’o v7 (30). Filtered reads from each sample were mapped against the associated assembly to provide an estimate of contig abundance using Bowtie 2 (24). Resulting SAM files were converted to BAM files before being sorted and indexed using SAMtools (25). Contig coverage depth was translated from each BAM file, before separately binning contigs >1000 bp with MaxBin v2.2.6 (31) and CONCOCT v1.1.0 (32) and binning contigs >1500 bp with MetaBAT 2 v2.12.1 (33).

DAS Tool was applied to the output from all three bin predictors, generating a catalogue of 196 bins from five samples (34). All bins were profiled against the BAM file for their source metagenomic sample using the anvi’o ‘anvi-profile’ workflow (30). Using the ‘anvi-interactive’ tool, each bin was refined manually according to GC content, single copy core gene (SCG) taxonomy and coverage as well as detection statistics. CheckM v1.0.11 (35) was used for quality assessment of all bins using the lineage_wf function. Bins showing >50% completion and <10% contamination were assessed for quality score (defined as estimated genome completeness score minus five times estimated contamination score) (36). Bins with <70% completion and/or a quality score of <50 were categorised as low-quality metagenome-assembled genomes (MAGs) (n=29); those with >70% completion, <10% contamination and quality score >50 were categorised as medium-quality MAGs (n=68) and those with >90% completion, <5% contamination and quality score >50 were classified as high-quality MAGs (n=55).

### Taxonomic and phylogenetic profiling of MAGs

Medium- and high-quality MAGs from all five samples were de-replicated at 95% average nucleotide identity (ANI) with a default aligned fraction of >10% using dRep v2.0 (37), to create a non-redundant species catalogue. Clustering at 99% ANI was used to identify a non-redundant strain catalogue and select a representative MAG per strain. CompareM v0.1.1 (38) was used to assign Average Amino-acid Identity (AAI) values followed by AAI clustering at 60% to allow delineation at the genus level.

The Genome Taxonomy Database Toolkit (GTDB-Tk) v1.4.1 (39), the Contig Annotation Tool (CAT/BAT) v5.2.3 (40) and ReferenceSeeker v1.4 (41) were used to perform taxonomic assignment of representative MAGs at strain-level compared to the ‘GTDB release 95’, ‘NCBI nr (2021-01-07)’ and ‘NCBI RefSeq release 201’ databases, respectively. Where taxonomic assignments differed between GTDB-Tk, CAT/BAT or ReferenceSeeker, GTDB-Tk assignments took precedence. Only when no species-level GTDB taxonomy was available did we adopt assignments according to CAT/BAT or ReferenceSeeker (11% of assignments). Phylogeny for our final de-replicated catalogue of MAGs was performed by aligning and concatenating a set of sixteen ribosomal protein sequences (ribosomal proteins L1, L2, L3, L4, L5, L6, L14, L16, L18, L22, L24, S3, S8, S10, S17 and S19), an approach previously used to reconstruct the tree of life (42). Ribosomal sequences were extracted using anvi’o before alignment using MUSCLE v3.8.155 (43) and refinement using trimAl v1.4 (44). A maximum-likelihood tree was constructed using FastTree v2.1 (45). All novel metagenomic species were confirmed as monophyletic, drawing on all publicly available genomes from the genus to which they had been assigned by GTDB genus (with genomes retrieved from NCBI). Proteomes were predicted using Prodigal v2.6.1 (46) before comparison against 400 universal marker proteins using PhyloPhlAn v3.0.58 (47) in accordance with diamond v0.9.34 (48). Multiple sequence alignment and subsequent refinement was performed using MAFFT v7.271 (49) and trimAl v1.4 (44) before tree construction using FastTree v2.1 and RAxML v8.2.12 (45, 50). All trees were subsequently visualised and manually annotated using iTol v5.7.

### Abundance and metabolic profiling of MAGs

To estimate the proportion of reads within each BioSample represented by our final, de- replicated MAG catalogue, contigs from the non-redundant MAG catalogue were concatenated and filtered reads aligned back to this MAG database using Bowtie 2 (24). Ordered BAM files were assessed using anvi’o (35) to calculate coverage statistics per-contig, allowing the calculation of mean coverage across each assembled genome. Species distribution analyses were conducted using the Vegan package in R (51) before visualisation using ggplot2 (52).

Functional profiling of high- and medium-quality MAGs (n=123) was performed using DRAM (Distilled and Refined Annotation of Metabolism) at a minimum contig length of 1000bp (53). Predicted amino-acid sequences identified by Prodigal in metagenome mode (46) were searched against KOfam, Pfam, and CAZy databases. tRNA and rRNA sequences were identified in MAGs using tRNAscan-SE (54) and Barrnap v0.9 (55), respectively.

### Bacteriophage identification and characterisation

VirSorter v1.0.5 (56) was applied to all contigs >5kb within each BioSample. Contig sequences classified by VirSorter as Category 1 (“most confident”) or Category 2 (“likely”) were considered for further analysis. Candidate bacteriophage sequences were assessed for completeness and contamination, using CheckV v0.7.0 (57), retaining only the sequences classified as “High-quality” (>90% completeness) or “complete”. These sequences were collated and de-replicated using rapid genome pairwise clustering at 95% ANI with an aligned fraction of ≥ 70% to generate a catalogue of bacteriophage genome sequences. For dereplication clustering, all-vs-all genome comparisons were performed using BLASTn before ANI based clustering using the ‘anicalc’ and ‘aniclust’ CheckV scripts sequentially.

Bacteriophage contigs from the catalogue were used as queries in a BLASTn search against the NCBI non-redundant nucleotide database (conducted on 21/12/2020) using an e-value of ≤1e-5. Only matches with a query cover >50% and percentage ID >70% were selected as being significant. Initial taxonomic classification of phage genomes at order and family level was performed using Demovir (58) against a viral subset of non-redundant TrEMBL database with an e-value of ≤1e-5. For each viral contig, individual coding sequences were predicted using Prodigal (46), before concatenation for input into vCONTACT2 v0.9.19 (59) for construction of a gene-sharing network incorporating a de- replicated RefSeq database of reference prokaryotic virus genomes. The resulting network was visualised using Cystoscape v3.8.0 (60).

## Results

### Reference-based profiling documents microbial diversity

Whole genome sequencing of five faecal samples derived from 12-month-old Thoroughbred horses, each yielded >6 ng/µl DNA and collectively generated >280 million paired reads or >84 Gbp of sequence data. Reads derived from the horse genome accounted for <1% of reads from each sample (S1 Table). We initially analysed reads using the k-mer-based program Kraken 2, followed by refined phylogenetic analysis via the allied program Bracken. Such analyses revealed unexpected novelty and diversity in the equine faecal microbiome, with >65% of sequence reads in each sample classified by Kraken as “unassigned”, i.e. from unknown organisms (S2 Table). Assignable reads represented all three domains of life, as well as viruses, although bacteria predominated, accounting for >95% of assigned reads.

Bacterial reads were predominantly assigned to the four phyla in the NCBI taxonomy most commonly associated with animal gut microbiomes—Proteobacteria, Firmicutes, Bacteroidetes and Actinobacteria. However, the Kraken 2 profiles also provided evidence of over thirty additional bacterial phyla in this ecosystem. Many of these appear to be novel in the context of the horse gut, including *Deinococcus-Thermus*, *Thermotogae* and the *Candidatus* phylum Cloacimonetes (also called WWE1), which has been reported almost exclusively from anaerobic fermenters and the aqueous environment (61, 62). However, as this phylum has recently been detected in soil fertilised with manure from dairy cattle, chickens and swine and has been implicated in anaerobic digestion of cellulose, it may play important similar roles in the vertebrate gut (62, 63). Interestingly, in four of the five samples more than a thousand reads were assigned to *Candidatus* Saccharibacteria, a phylum from the candidate phyla radiation, which is home to bacteria that live as bacterial epibionts (64). Reads assigned to eukaryotes provided evidence of budding yeasts and apicoplexan parasites in these samples.

Remarkably, two samples show a very high relative abundance of reads assigned to the genus *Acinetobacter* (20% and 9.4% of all reads or 57% and 34% of classified reads), mirroring similar findings on two healthy horses in a previous study using 16S rRNA gene sequences (65). Bracken assigns these reads to an implausible thirty-one species of *Acinetobacter*, which is more likely to represent misassignment of reads rather than genuine diversity within this genus in this context.

### Over a hundred newly named bacterial species

We generated almost 200 non-redundant bins from single-sample assemblies using three different approaches to binning. 123 bins represent medium- or high-quality metagenome-assembled genomes (MAGs), 96 with ≥15 amino acid tRNAs (S3 and S4 Tables). Genome sizes ranged from ∼0.5 to 3.8 Mbp, while GC content ranged from 31% to 60%. De- replication at 95% ANI clustered MAGs into 110 metagenomic species, spanning ten phyla (Fig 1a). According to GTDB, around half (48%) of the metagenomic species belonged to the *Bacteroidota*, while just over a third (35%) belonged to the *Firmicutes* (split by GTDB into *Firmicutes, Firmicutes_A* and *Firmicutes_C*). Only twelve species of bacterial species from the horse gut had been previously defined and delineated: eight with validly published Latin binomials and four simply with alphanumerical designations assigned by GTDB (S5 Table).

**Fig 1.**
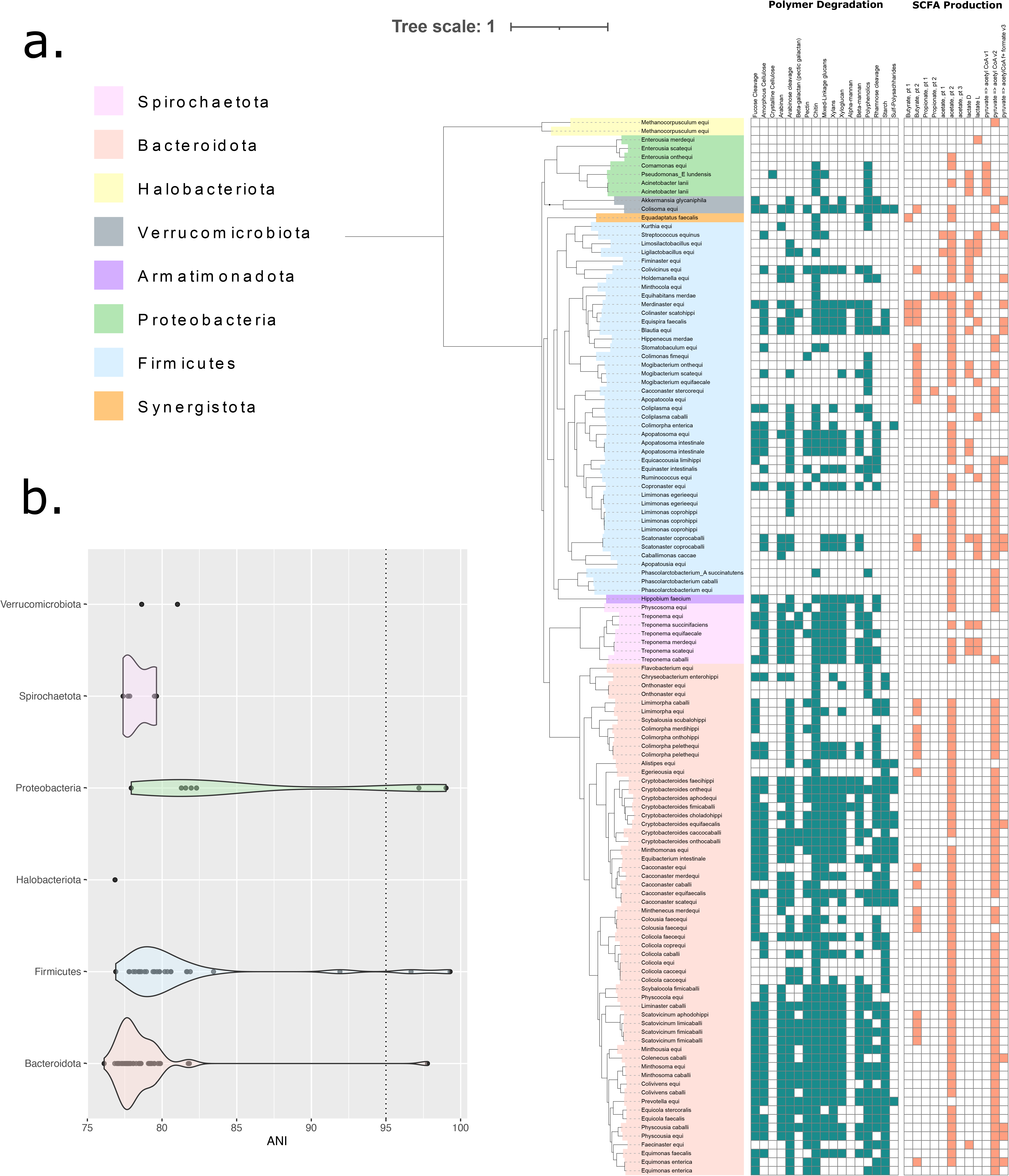
Taxonomic classification of 110 Metagenomic species derived from 5 metagenomic equine faecal samples. (**A**) Depicted as a phylogenetic tree - where phylum, as assigned by GTDB, is indicated by colour range. The tree was based upon an alignment of 16 concatenated ribosomal proteins and constructed using FastTree. The final tree was visualised and manually annotated using the online iTOLv5.7 tool. Phylum-level taxonomy is described by branch colour according to GTDB designation (Phyla with an alphabetical suffix have been collapsed). The presence (blue filled) or absence (hollow) of genes associated with catalysing carbohydrate degradation (blue) or aiding in the metabolism of short chain fatty acids (red) are reported in the associated binary plot. Hemicellulose substrates have been clustered and highlighted in red text. (**B)** Average Nucleotide Identity (ANI) between recovered MAGs and their closest representative within the GTDB database (release 95). Only MAGs placed within a previously recognised genus, and whereby this taxonomic assignment was inclusive of an ANI measurement, are shown. Individual plots are coloured according to GTDB designated phylum, with phyla assigned an alphabetical suffix being collapsed. A dotted line is placed at 95% ANI, representing the utilised species-level boundary.

Two of the species with validly published names, *Ligilactobacillus hayakitensis* (synonym *Lactobacillus hayakitensis*) (66) and *Limosilactobacillus equigenerosi* (synonym *Lactobacillus equigenerosi*) (67), have been previously cultured from the faeces of thoroughbred racehorses and are thought to be positively associated with equine intestinal health (68). Similarly, the species *Streptococcus equinus* was named in the early twentieth century after its association with horse dung and has been repeatedly isolated from this source (69, 70). Another of the named species found among our MAGs, *Treponema succinifaciens,* has been reported from the equine gut by 16S studies (71), but ours represents the first report of a genome from this species in this setting.

The recently named species *Acinetobacter lanii* (72) has been isolated from the Tibetan wild ass *Equus kiang*, but our MAG represents the first report of an association between this species and the domesticated horse. Similarly, although the genus *Phascolarctobacterium* is known to inhabit the horse gut (15, 16), here we provide the first evidence of a specific link between the horse and the species *P. succinatutens*, previously found in human and pig faeces (73, 74). Our metagenomic species provide the first report in the horse of the species *Pseudomonas lundensis* (first isolated from meat, but now recognised as an emerging pathogen of humans (75, 76)) and *of Akkermansia glycaniphila*, previously isolated from the faeces of the reticulated python (77).

Among our metagenomic species, eighty-five represent new candidate species within fifty bacterial genera previously delineated by GTDB (Fig 1b). All fifty of these genera occur in the gut microbiota of at least one additional mammalian host species. Twelve of our metagenomic species that could be assigned only to the level of family fell into ten clusters (delineated at 60% AAI) representing novel candidate genera from seven different families (S6 Table). The archaeal genus *Methanocorpusculum* is thought to play a role in methane production in the equine gut (78). Here, we have delineated a novel species from this ecosystem: Candidatus *Methanocorpusculum equi*.

Building on our recent efforts with the chicken gut microbiome and with the automated creation of well-formed Latin names, we have created *Candidatus* names (abbreviated as *Ca.*) for all the unnamed taxa revealed by our metagenomic analyses (Table 1). We also created Latin names for species and genera recognised by GTDB, but previously assigned only alphanumeric designations. For taxa found only in the horse, we created names that incorporated Greek or Latin roots for this host (e.g. *Ca.* Equimonas). However, if searches of the GTDB and NCBI databases suggested that genera had representatives in other gut microbiomes, we opted for names that specified gut or faeces as habitat (e.g. *Ca.* Limimonas).

**Table 1. Protologues for newly named *Candidatus* genera and species.**

Protologues for new *Candidatus* taxa identified by analysis of metagenome-assembled genomes from equine faeces.

### Description of *Candidatus* Alistipes equi sp. nov

*Candidatus* Alistipes equi (e’qui. L. gen. masc. n. *equi*, of a horse) A bacterial species identified by metagenomic analyses. This species includes all bacteria with genomes that show ≥95% average nucleotide identity (ANI) to the type genome for the species to which we have assigned the MAG ID E3_MB2_80 and which is available via NCBI BioSample SAMN18472495. The GC content of the type genome is 40.8 % and the genome length is 2.08 Mbp.

### Description of *Candidatus* Apopatocola gen. nov

*Candidatus* Apopatocola (A.po.pa.to’cola. Gr. masc. n. *apopatos*, dung; N.L. masc./fem. suffix *–cola*, an inhabitant; N.L. fem. n. *Apopatocola* a microbe associated with faeces) A bacterial genus identified by metagenomic analyses. The genus includes all bacteria with genomes that show ≥60% average amino acid identity (AAI) to the genome of the type strain from the type species *Candidatus* Apopatocola equi. This is a new name for the GTDB alphanumeric genus UBA738, which is found in diverse mammalian guts. This genus has been assigned by GTDB-Tk v1.3.0 working on GTDB Release 05-RS95 (39, 79) to the order *Oscillospirales* and to the family *Oscillospiraceae*.

### Description of *Candidatus* Apopatocola equi sp. nov

*Candidatus* Apopatocola equi (e’qui. L. gen. masc. n. *equi*, of a horse)

A bacterial species identified by metagenomic analyses. This species includes all bacteria with genomes that show ≥95% average nucleotide identity (ANI) to the type genome for the species to which we have assigned the MAG ID E1_MB2_75 and which is available via NCBI BioSample SAMN18472466. The GC content of the type genome is 59.6 % and the genome length is 1.56 Mbp.

### Description of *Candidatus* Apopatosoma gen. nov

*Candidatus* Apopatosoma (A.po.pa.to.so’ma. Gr. masc. n. *apopatos*, dung; Gr. neut. n. *soma*, a body; N.L. neut. n. *Apopatosoma*, a microbe associated with faeces)

A bacterial genus identified by metagenomic analyses. The genus includes all bacteria with genomes that show ≥60% average amino acid identity (AAI) to the genome of the type strain from the type species *Candidatus* Apopatosoma equi. This is a new name for the GTDB alphanumeric genus CAG-724, which is found in diverse mammalian guts. This genus has been assigned by GTDB-Tk v1.3.0 working on GTDB Release 05-RS95 (39, 79) to the order *Oscillospirales* and to the family *CAG-272*.

### Description of *Candidatus* Apopatosoma equi sp. nov

*Candidatus* Apopatosoma equi (e’qui. L. gen. masc. n. *equi*, of a horse)

A bacterial species identified by metagenomic analyses. This species includes all bacteria with genomes that show ≥95% average nucleotide identity (ANI) to the type genome for the species to which we have assigned the MAG ID E2_100 and which is available via NCBI BioSample SAMN18472471. The GC content of the type genome is 49.5 % and the genome length is 1.54 Mbp.

### Description of *Candidatus* Apopatosoma intestinale sp. nov

*Candidatus* Apopatosoma intestinale (in.tes.ti.na’le. N.L. neut. adj. *intestinale*, pertaining to the intestines)

A bacterial species identified by metagenomic analyses. This species includes all bacteria with genomes that show ≥95% average nucleotide identity (ANI) to the type genome for the species to which we have assigned the MAG ID E5_133 and which is available via NCBI BioSample SAMN18472535. This is a new name for the alphanumeric GTDB species sp003524145, which is found in diverse mammalian guts. The GC content of the type genome is 53.8 % and the genome length is 1.55 Mbp.

### Description of *Candidatus* Apopatousia gen. nov

*Candidatus* Apopatousia (A.po.pat.ou’s.ia. Gr. masc. n. *apopatos*, dung; Gr. fem. n. *ousia*, an essence; N.L. fem. n. *Apopatousia*, a microbe associated with faeces)

A bacterial genus identified by metagenomic analyses. The genus includes all bacteria with genomes that show ≥60% average amino acid identity (AAI) to the genome of the type strain from the type species *Candidatus* Apopatousia equi. This is a new name for the GTDB alphanumeric genus UBA9845, which is found in diverse mammalian guts. This genus has been assigned by GTDB-Tk v1.3.0 working on GTDB Release 05-RS95 (39, 79) to the order *Christensenellales* and to the family *UBA1242*.

### Description of *Candidatus* Apopatousia equi sp. nov

*Candidatus* Apopatousia equi (e’qui. L. gen. masc. n. *equi*, of a horse)

A bacterial species identified by metagenomic analyses. This species includes all bacteria with genomes that show ≥95% average nucleotide identity (ANI) to the type genome for the species to which we have assigned the MAG ID E5_MB2_6 and which is available via NCBI BioSample SAMN18472550. The GC content of the type genome is 31.9 % and the genome length is 0.57 Mbp.

### Description of *Candidatus* Blautia equi sp. nov

*Candidatus* Blautia equi (e’qui. L. gen. masc. n. *equi*, of a horse)

A bacterial species identified by metagenomic analyses. This species includes all bacteria with genomes that show ≥95% average nucleotide identity (ANI) to the type genome for the species to which we have assigned the MAG ID E4_MB2_89 and which is available via NCBI BioSample SAMN18472531. GTDB has assigned. This species to a genus marked with an alphabetical suffix. However, as this genus designation cannot be incorporated into a well-formed binomial, in naming. This species, we have used the current validly published name for the genus. The GC content of the type genome is 48 % and the genome length is 2.14 Mbp.

### Description of *Candidatus* Caballimonas gen. nov

*Candidatus* Caballimonas (Ca.bal.li.mo’nas. L. masc. n. *caballus*, a horse; L. fem. n. *monas*, a monad; N.L. fem. n. *Caballimonas*, a microbe associated with horses)

A bacterial genus identified by metagenomic analyses. The genus includes all bacteria with genomes that show ≥60% average amino acid identity (AAI) to the genome of the type strain from the type species *Candidatus* Caballimonas caccae. This genus has been assigned by GTDB-Tk v1.3.0 working on GTDB Release 05-RS95 (39, 79) to the order *Christensenellales* and to the family *Borkfalkiaceae*.

### Description of *Candidatus* Caballimonas caccae sp. nov

*Candidatus* Caballimonas caccae (cac’cae. Gr. fem. n. *kakke*, faeces; N.L. gen. n. *caccae*, of faeces)

A bacterial species identified by metagenomic analyses. This species includes all bacteria with genomes that show ≥95% average nucleotide identity (ANI) to the type genome for the species to which we have assigned the MAG ID E3_31 and which is available via NCBI BioSample SAMN18472486. The GC content of the type genome is 34.9 % and the genome length is 0.91 Mbp.

### Description of *Candidatus* Cacconaster gen. nov

*Candidatus* Cacconaster (Cac.co.nas’ter. Gr. fem. n. *kakke*, dung; Gr. masc. n. *naster*, an inhabitant; N.L. masc. n. *Cacconaster*, a microbe associated with faeces)

A bacterial genus identified by metagenomic analyses. The genus includes all bacteria with genomes that show ≥60% average amino acid identity (AAI) to the genome of the type strain from the type species *Candidatus* Cacconaster caballi. This is a new name for the GTDB alphanumeric genus Bact-11, which is found in diverse mammalian guts. This genus has been assigned by GTDB-Tk v1.3.0 working on GTDB Release 05-RS95 (39, 79) to the order *Bacteroidales* and to the family *UBA932*,

### Description of *Candidatus* Cacconaster caballi sp. nov

*Candidatus* Cacconaster caballi (ca.bal’li. L. gen. masc. n. *caballi*, of a horse)

A bacterial species identified by metagenomic analyses. This species includes all bacteria with genomes that show ≥95% average nucleotide identity (ANI) to the type genome for the species to which we have assigned the MAG ID E2_MB2_69 and which is available via NCBI BioSample SAMN18472478. The GC content of the type genome is 50.7 % and the genome length is 1.38 Mbp.

### Description of *Candidatus* Cacconaster equi sp. nov

*Candidatus* Cacconaster equi (e’qui. L. gen. masc. n. *equi*, of a horse)

A bacterial species identified by metagenomic analyses. This species includes all bacteria with genomes that show ≥95% average nucleotide identity (ANI) to the type genome for the species to which we have assigned the MAG ID E1_MB2_89 and which is available via NCBI BioSample SAMN18472469. The GC content of the type genome is 48.5 % and the genome length is 1.65 Mbp.

### Description of *Candidatus* Cacconaster equifaecalis sp. nov

*Candidatus* Cacconaster equifaecalis (e.qui.fae.ca’lis. L. masc. n. *equus*, a horse; N.L. masc. adj. *faecalis*, faecal; N.L. masc. adj. *equifaecalis,* associated with the faeces of horses)

A bacterial species identified by metagenomic analyses. This species includes all bacteria with genomes that show ≥95% average nucleotide identity (ANI) to the type genome for the species to which we have assigned the MAG ID E5_MB2_108 and which is available via NCBI BioSample SAMN18472541. The GC content of the type genome is 51.7 % and the genome length is 1.71 Mbp.

### Description of *Candidatus* Cacconaster merdequi sp. nov

*Candidatus* Cacconaster merdequi (merd.e’qui. L. fem. n. *merda*, faeces; L. masc. n. *equus*, a horse; N.L. gen. n. *merdequi*, associated with the faeces of horses)

A bacterial species identified by metagenomic analyses. This species includes all bacteria with genomes that show ≥95% average nucleotide identity (ANI) to the type genome for the species to which we have assigned the MAG ID E5_MB2_33 and which is available via NCBI BioSample SAMN18472547. The GC content of the type genome is 49 % and the genome length is 1.90 Mbp.

### Description of *Candidatus* Cacconaster scatequi sp. nov

*Candidatus* Cacconaster scatequi (scat.e’qui. Gr. neut. n. *skor*, *skatos*, dung; L. masc. n. *equus*, a horse; N.L. gen. n. *scatequi*, associated with the faeces of horses)

A bacterial species identified by metagenomic analyses. This species includes all bacteria with genomes that show ≥95% average nucleotide identity (ANI) to the type genome for the species to which we have assigned the MAG ID E3_MB2_97 and which is available via NCBI BioSample SAMN18472499. The GC content of the type genome is 50.6 % and the genome length is 1.90 Mbp.

### Description of *Candidatus* Cacconaster stercorequi sp. nov

*Candidatus* Cacconaster stercorequi (ster.cor.e’qui. L. masc. n. *stercus*, *stercoris*, dung; L. masc. n. *equus*, a horse; N.L. gen. n. *stercorequi*, associated with the faeces of horses)

A bacterial species identified by metagenomic analyses. This species includes all bacteria with genomes that show ≥95% average nucleotide identity (ANI) to the type genome for the species to which we have assigned the MAG ID E4_MB2_17 and which is available via NCBI BioSample SAMN18472518. The GC content of the type genome is 54.5 % and the genome length is 1.83 Mbp.

### Description of *Candidatus* Chryseobacterium enterohippi sp. nov

*Candidatus* Chryseobacterium enterohippi (en.te.ro.hip’pi. Gr. neut. n. *enteron*, gut, bowel, intestine; Gr. masc./fem. n. *hippos,* a horse; N.L. gen. n. *enterohippi*, associated with the horse gut)

A bacterial species identified by metagenomic analyses. This species includes all bacteria with genomes that show ≥95% average nucleotide identity (ANI) to the type genome for the species to which we have assigned the MAG ID E1_189 and which is available via NCBI BioSample SAMN18472455. The GC content of the type genome is 34.3 % and the genome length is 2.05 Mbp.

### Description of *Candidatus* Colenecus gen. nov

*Candidatus* Colenecus (Col.en.e’cus. L. neut. n. *colon*, large intestine; N.L. masc. n. *enecus*, an inhabitant; N.L. masc. n. *Colenecus*, a microbe associated with the large intestine)

A bacterial genus identified by metagenomic analyses. The genus includes all bacteria with genomes that show ≥60% average amino acid identity (AAI) to the genome of the type strain from the type species *Candidatus* Colenecus caballi. This is a new name for the GTDB alphanumeric genus UBA1179, which is found in diverse mammalian guts. This genus has been assigned by GTDB-Tk v1.3.0 working on GTDB Release 05-RS95 (39, 79) to the order *Bacteroidales* and to the family *Bacteroidaceae*.

### Description of *Candidatus* Colenecus caballi sp. nov

*Candidatus* Colenecus caballi (ca.bal’li. L. gen. masc. n. *caballi*, of a horse)

A bacterial species identified by metagenomic analyses. This species includes all bacteria with genomes that show ≥95% average nucleotide identity (ANI) to the type genome for the species to which we have assigned the MAG ID E3_160 and which is available via NCBI BioSample SAMN18472483. The GC content of the type genome is 49.7 % and the genome length is 2.25 Mbp.

### Description of *Candidatus* Colicola gen. nov

*Candidatus* Colicola (Co.li.co’la. L. neut. n. *colon*, large intestine; N.L. masc./fem. suffix *– cola*, an inhabitant; N.L. fem. n. *Colicola*, a microbe associated with the large intestine)

A bacterial genus identified by metagenomic analyses. The genus includes all bacteria with genomes that show ≥60% average amino acid identity (AAI) to the genome of the type strain from the type species *Candidatus* Colicola caballi. This is a new name for the GTDB alphanumeric genus RF16, which is found in diverse mammalian guts. This genus has been assigned by GTDB-Tk v1.3.0 working on GTDB Release 05-RS95 (39, 79) to the order *Bacteroidales* and to the family *Paludibacteraceae*.

### Description of *Candidatus* Colicola caballi sp. nov

*Candidatus* Colicola caballi (ca.bal’li. L. gen. masc. n. *caballi*, of a horse)

A bacterial species identified by metagenomic analyses. This species includes all bacteria with genomes that show ≥95% average nucleotide identity (ANI) to the type genome for the species to which we have assigned the MAG ID E1_MB2_58 and which is available via NCBI BioSample SAMN18472465. The GC content of the type genome is 46.6 % and the genome length is 1.45 Mbp.

### Description of *Candidatus* Colicola caccequi sp. nov

*Candidatus* Colicola caccequi (cacc.e’qui. Gr. fem. n. *kakke*, faeces; L. masc. n. *equus*, a horse; N.L. gen. n. *caccequi*, associated with the faeces of horses)

A bacterial species identified by metagenomic analyses. This species includes all bacteria with genomes that show ≥95% average nucleotide identity (ANI) to the type genome for the species to which we have assigned the MAG ID E4_134 and which is available via NCBI BioSample SAMN18472502. The GC content of the type genome is 44.2 % and the genome length is 1.71 Mbp.

### Description of *Candidatus* Colicola coprequi sp. nov

*Candidatus* Colicola coprequi (copr.e’qui. Gr. fem. n. *kopros*, dung; L. masc. n. *equus*, a horse; N.L. gen. n. *coprequi*, associated with the faeces of horses)

A bacterial species identified by metagenomic analyses. This species includes all bacteria with genomes that show ≥95% average nucleotide identity (ANI) to the type genome for the species to which we have assigned the MAG ID E2_MB2_30 and which is available via NCBI BioSample SAMN18472476. The GC content of the type genome is 46.1 % and the genome length is 1.53 Mbp.

### Description of *Candidatus* Colicola equi sp. nov

*Candidatus* Colicola equi (e’qui. L. gen. masc. n. *equi*, of a horse)

A bacterial species identified by metagenomic analyses. This species includes all bacteria with genomes that show ≥95% average nucleotide identity (ANI) to the type genome for the species to which we have assigned the MAG ID E1_186 and which is available via NCBI BioSample SAMN18472454. The GC content of the type genome is 44.4 % and the genome length is 2.05 Mbp.

### Description of *Candidatus* Colicola faecequi sp. nov

*Candidatus* Colicola faecequi (faec.e’qui. L. fem. n. *faex*, *faeces*, *dregs;* L. masc. n. *equus*, a horse; N.L. gen. n. *faecequi*, associated with the faeces of horses)

A bacterial species identified by metagenomic analyses. This species includes all bacteria with genomes that show ≥95% average nucleotide identity (ANI) to the type genome for the species to which we have assigned the MAG ID E4_MB2_124 and which is available via NCBI BioSample SAMN18472515. The GC content of the type genome is 52.3 % and the genome length is 1.86 Mbp.

### Description of *Candidatus* Colimonas gen. nov

*Candidatus* Colimonas (Co.li.mo’nas. L. neut. n. *colon*, large intestine; L. fem. n. *monas*, a monad; N.L. fem. n. *Colimonas*, a microbe associated with the large intestine)

A bacterial genus identified by metagenomic analyses. The genus includes all bacteria with genomes that show ≥60% average amino acid identity (AAI) to the genome of the type strain from the type species *Candidatus* Colimonas fimequi. This is a new name for the GTDB alphanumeric genus UBA1191, which is found in diverse mammalian guts. This genus has been assigned by GTDB-Tk v1.3.0 working on GTDB Release 05-RS95 (39, 79) to the order *Peptostreptococcales* and to the family *Anaerovoracaceae*.

### Description of *Candidatus* Colimonas fimequi sp. nov

*Candidatus* Colimonas fimequi (fim.e’qui. L. masc. n. *fimus*, dung; L. masc. n. *equus*, a horse; N.L. gen. n. *fimequi*, associated with the faeces of horses)

A bacterial species identified by metagenomic analyses. This species includes all bacteria with genomes that show ≥95% average nucleotide identity (ANI) to the type genome for the species to which we have assigned the MAG ID E4_13 and which is available via NCBI BioSample SAMN18472501. The GC content of the type genome is 44.3 % and the genome length is 1.70 Mbp.

### Description of *Candidatus* Colimorpha gen. nov

*Candidatus* Colimorpha (Co.li.mor’pha. L. neut. n. *colon*, large intestine; Gr. fem. n. *morphe*, a form, shape; N.L. fem. n. *Colimorpha*, a microbe associated with the large intestine)

A bacterial genus identified by metagenomic analyses. The genus includes all bacteria with genomes that show ≥60% average amino acid identity (AAI) to the genome of the type strain from the type species *Candidatus* Colimorpha merdihippi. This is a new name for the GTDB alphanumeric genus UBA1711, which is found in diverse mammalian guts. This genus has been assigned by GTDB-Tk v1.3.0 working on GTDB Release 05-RS95 (39, 79) to the order *Bacteroidales* and to the family *P3*.

### Description of *Candidatus* Colimorpha enterica sp. nov

*Candidatus* Colimorpha enterica (en.te’ri.ca. Gr. neut. n. *enteron*, gut, bowel, intestine; L. fem. adj. suff. *-ica*, pertaining to; N.L. fem. adj. *enterica*, pertaining to intestine)

A bacterial species identified by metagenomic analyses. This species includes all bacteria with genomes that show ≥95% average nucleotide identity (ANI) to the type genome for the species to which we have assigned the MAG ID E3_60 and which is available via NCBI BioSample SAMN18472488. This is a new name for the alphanumeric GTDB species sp000433515, which is found in diverse mammalian guts. The GC content of the type genome is 52.3 % and the genome length is 1.43 Mbp.

### Description of *Candidatus* Colimorpha merdihippi sp. nov

*Candidatus* Colimorpha merdihippi (mer.di.hip’pi. L. fem. n. *merda*, faeces; Gr. masc./fem. n. *hippos*, a horse; N.L. gen. n. *merdihippi*, associated with the faeces of horses)

A bacterial species identified by metagenomic analyses. This species includes all bacteria with genomes that show ≥95% average nucleotide identity (ANI) to the type genome for the species to which we have assigned the MAG ID E1_90 and which is available via NCBI BioSample SAMN18472457. The GC content of the type genome is 48.5 % and the genome length is 3.11 Mbp.

### Description of *Candidatus* Colimorpha onthohippi sp. nov

*Candidatus* Colimorpha onthohippi (on.tho.hip’pi. Gr. masc. n. *onthos*, dung; Gr. masc./fem. n. *hippos*, a horse; N.L. gen. n. *onthohippi*, associated with the faeces of horses)

A bacterial species identified by metagenomic analyses. This species includes all bacteria with genomes that show ≥95% average nucleotide identity (ANI) to the type genome for the species to which we have assigned the MAG ID E5_36 and which is available via NCBI

BioSample SAMN18472537. The GC content of the type genome is 46.2 % and the genome length is 2.04 Mbp.

### Description of *Candidatus* Colimorpha pelethequi sp. nov

*Candidatus* Colimorpha pelethequi (pe.leth.e’qui. Gr. masc. n. *pelethos*, dung; L. masc. n. *equus*, a horse; N.L. gen. n. *pelethequi*, associated with the faeces of horses)

A bacterial species identified by metagenomic analyses. This species includes all bacteria with genomes that show ≥95% average nucleotide identity (ANI) to the type genome for the species to which we have assigned the MAG ID E5_MB2_81 and which is available via NCBI BioSample SAMN18472551. The GC content of the type genome is 46.7 % and the genome length is 2.38 Mbp.

### Description of *Candidatus* Colinaster gen. nov

*Candidatus* Colinaster (Co.li.nas’ter. L. neut. n. *colon*, large intestine; Gr. masc. n. *naster*, an inhabitant; N.L. masc. n. *Colinaster* a microbe associated with the large intestine)

A bacterial genus identified by metagenomic analyses. The genus includes all bacteria with genomes that show ≥60% average amino acid identity (AAI) to the genome of the type strain from the type species *Candidatus* Colinaster scatohippi. This is a new name for the GTDB alphanumeric genus UBA1712, which is found in diverse mammalian guts. This genus has been assigned by GTDB-Tk v1.3.0 working on GTDB Release 05-RS95 (39, 79) to the order *Lachnospirales* and to the family *Lachnospiraceae*.

### Description of *Candidatus* Colinaster scatohippi sp. nov

*Candidatus* Colinaster scatohippi (sca.to.hip’pi. Gr. neut. n. *skor*, *skatos*, dung; Gr. masc./fem. n. *hippos*a horse; N.L. gen. n. *scatohippi*, associated with the faeces of horses) A bacterial species identified by metagenomic analyses. This species includes all bacteria with genomes that show ≥95% average nucleotide identity (ANI) to the type genome for the species to which we have assigned the MAG ID E4_MB2_45 and which is available via NCBI BioSample SAMN18472524. The GC content of the type genome is 38.7 % and the genome length is 2.18 Mbp.

### Description of *Candidatus* Coliplasma gen. nov

*Candidatus* Coliplasma (Co.li.plas’ma. L. neut. n. *colon*, large intestine; Gr. neut. n. *plasma*, a form; N.L. neut. n. *Coliplasma*, a microbe associated with the large intestine)

A bacterial genus identified by metagenomic analyses. The genus includes all bacteria with genomes that show ≥60% average amino acid identity (AAI) to the genome of the type strain from the type species *Candidatus* Coliplasma caballi. This is a new name for the GTDB alphanumeric genus UBA1752, which is found in diverse mammalian guts. This genus has been assigned by GTDB-Tk v1.3.0 working on GTDB Release 05-RS95 (39, 79) to the order *Oscillospirales* and to the family *CAG-382*.

### Description of *Candidatus* Coliplasma caballi sp. nov

*Candidatus* Coliplasma caballi (ca.bal’li. L. gen. masc. n. *caballi*, of a horse)

A bacterial species identified by metagenomic analyses. This species includes all bacteria with genomes that show ≥95% average nucleotide identity (ANI) to the type genome for the species to which we have assigned the MAG ID E3_MB2_28 and which is available via NCBI BioSample SAMN18472492. The GC content of the type genome is 54.8 % and the genome length is 1.41 Mbp.

### Description of *Candidatus* Coliplasma equi sp. nov

*Candidatus* Coliplasma equi (e’qui. L. gen. masc. n. *equi*, of a horse)

A bacterial species identified by metagenomic analyses. This species includes all bacteria with genomes that show ≥95% average nucleotide identity (ANI) to the type genome for the species to which we have assigned the MAG ID E3_142 and which is available via NCBI BioSample SAMN18472481. The GC content of the type genome is 49.7 % and the genome length is 1.52 Mbp.

### Description of *Candidatus* Colisoma gen. nov

*Candidatus* Colisoma (Co.li.so’ma. L. neut. n. *colon*, large intestine; Gr. neut. n. *soma*, a body; N.L. neut. n. *Colisoma*, a microbe associated with the large intestine)

A bacterial genus identified by metagenomic analyses. The genus includes all bacteria with genomes that show ≥60% average amino acid identity (AAI) to the genome of the type strain from the type species *Candidatus* Colisoma equi. This is a new name for the GTDB alphanumeric genus UBA1067, which is found in diverse mammalian guts. This genus has been assigned by GTDB-Tk v1.3.0 working on GTDB Release 05-RS95 (39, 79) to the order *RFP12* and to the family *UBA1067*.

### Description of *Candidatus* Colisoma equi sp. nov

*Candidatus* Colisoma equi (e’qui. L. gen. masc. n. *equi*, of a horse)

A bacterial species identified by metagenomic analyses. This species includes all bacteria with genomes that show ≥95% average nucleotide identity (ANI) to the type genome for the species to which we have assigned the MAG ID E4_MB2_14 and which is available via NCBI BioSample SAMN18472517. The GC content of the type genome is 60 % and the genome length is 2.52 Mbp.

### Description of *Candidatus* Colivicinus gen. nov

*Candidatus* Colivicinus (Co.li.vi’ci.nus. L. neut. n. *colon*, large intestine; N.L. masc. n. *vicinus*, a neighbour; N.L. masc. n. *Colivicinus*, a microbe associated with the large intestine) A bacterial genus identified by metagenomic analyses. The genus includes all bacteria with genomes that show ≥60% average amino acid identity (AAI) to the genome of the type strain from the type species *Candidatus* Colivicinus equi. This is a new name for the GTDB alphanumeric genus UBA636, which is found in diverse mammalian guts. This genus has been assigned by GTDB-Tk v1.3.0 working on GTDB Release 05-RS95 (39, 79) to the order *Erysipelotrichales* and to the family *Erysipelotrichaceae*.

### Description of *Candidatus* Colivicinus equi sp. nov

*Candidatus* Colivicinus equi (e’qui. L. gen. masc. n. *equi*, of a horse)

A bacterial species identified by metagenomic analyses. This species includes all bacteria with genomes that show ≥95% average nucleotide identity (ANI) to the type genome for the species to which we have assigned the MAG ID E4_MB2_36 and which is available via NCBI BioSample SAMN18472522. The GC content of the type genome is 31.9 % and the genome length is 1.69 Mbp.

### Description of *Candidatus* Colivivens gen. nov

*Candidatus* Colivivens (Co.li.vi’vens. L. neut. n. *colon*, large intestine; N.L. masc./fem. pres. part. *vivens*, living; N.L. fem. n. *Colivivens*, a microbe associated with the large intestine)

A bacterial genus identified by metagenomic analyses. The genus includes all bacteria with genomes that show ≥60% average amino acid identity (AAI) to the genome of the type strain from the type species *Candidatus* Colivivens caballi. This is a new name for the GTDB alphanumeric genus UBA1786, which is found in diverse mammalian guts. This genus has been assigned by GTDB-Tk v1.3.0 working on GTDB Release 05-RS95 (39, 79) to the order *Bacteroidales* and to the family *Bacteroidaceae*.

### Description of *Candidatus* Colivivens caballi sp. nov

*Candidatus* Colivivens caballi (ca.bal’li. L. gen. masc. n. *caballi*, of a horse)

A bacterial species identified by metagenomic analyses. This species includes all bacteria with genomes that show ≥95% average nucleotide identity (ANI) to the type genome for the species to which we have assigned the MAG ID E3_198 and which is available via NCBI BioSample SAMN18472484. The GC content of the type genome is 47.7 % and the genome length is 2.55 Mbp.

### Description of *Candidatus* Colivivens equi sp. nov

*Candidatus* Colivivens equi (e’qui. L. gen. masc. n. *equi*, of a horse)

A bacterial species identified by metagenomic analyses. This species includes all bacteria with genomes that show ≥95% average nucleotide identity (ANI) to the type genome for the species to which we have assigned the MAG ID E1_MB2_52 and which is available via NCBI BioSample SAMN18472463. The GC content of the type genome is 38.2 % and the genome length is 2.64 Mbp.

### Description of *Candidatus* Colousia gen. nov

*Candidatus* Colousia (Col.ou’s.ia. L. neut. n. *colon*, large intestine; Gr. fem. n. *ousia*, an essence; N.L. fem. n. *Colousia*, a microbe associated with the large intestine)

A bacterial genus identified by metagenomic analyses. The genus includes all bacteria with genomes that show ≥60% average amino acid identity (AAI) to the genome of the type strain from the type species *Candidatus* Colousia faecequi. This is a new name for the GTDB alphanumeric genus SFVR01, which is found in diverse mammalian guts. This genus has been assigned by GTDB-Tk v1.3.0 working on GTDB Release 05-RS95 (39, 79) to the order *Bacteroidales* and to the family *Paludibacteraceae*.

### Description of *Candidatus* Colousia faecequi sp. nov

*Candidatus* Colousia faecequi (faec.e’qui. L. fem. n. *faex*, *faeces*, *dregs*; L. masc. n. *equus*, a horse; N.L. gen. n. *faecequi,* associated with the faeces of horses)

A bacterial species identified by metagenomic analyses. This species includes all bacteria with genomes that show ≥95% average nucleotide identity (ANI) to the type genome for the species to which we have assigned the MAG ID E3_MB2_91 and which is available via NCBI BioSample SAMN18472498. The GC content of the type genome is 47.1 % and the genome length is 1.67 Mbp.

### Description of *Candidatus* Comamonas equi sp. nov

*Candidatus* Comamonas equi (e’qui. L. gen. masc. n. *equi*, of a horse)

A bacterial species identified by metagenomic analyses. This species includes all bacteria with genomes that show ≥95% average nucleotide identity (ANI) to the type genome for the species to which we have assigned the MAG ID E2_118 and which is available via NCBI BioSample SAMN18472472. The GC content of the type genome is 59.2 % and the genome length is 2.60 Mbp.

### Description of *Candidatus* Copronaster gen. nov

*Candidatus* Copronaster (Co.pro.nas’ter. Gr. fem. n. *kopros*, dung; Gr. masc. n. *naster*, an inhabitant; N.L. masc. n. *Copronaster*, a microbe associated with faeces)

A bacterial genus identified by metagenomic analyses. The genus includes all bacteria with genomes that show ≥60% average amino acid identity (AAI) to the genome of the type strain from the type species *Candidatus* Copronaster equi. This is a new name for the GTDB alphanumeric genus CAG-488, which is found in diverse mammalian guts. This genus has been assigned by GTDB-Tk v1.3.0 working on GTDB Release 05-RS95 (39, 79) to the order *Oscillospirales* and to the family *Acutalibacteraceae*.

### Description of *Candidatus* Copronaster equi sp. nov

*Candidatus* Copronaster equi (e’qui. L. gen. masc. n. *equi*, of a horse)

A bacterial species identified by metagenomic analyses. This species includes all bacteria with genomes that show ≥95% average nucleotide identity (ANI) to the type genome for the species to which we have assigned the MAG ID E5_MB2_59 and which is available via NCBI BioSample SAMN18472549. The GC content of the type genome is 39.1 % and the genome length is 1.98 Mbp.

### Description of *Candidatus* Cryptobacteroides aphodequi sp. nov

*Candidatus* Cryptobacteroides aphodequi (aph.od.e’qui. Gr. fem. n. *aphodo*s, dung; L. masc. n. *equus*, a horse; N.L. gen. n. *aphodequi*, associated with the faeces of horses)

A bacterial species identified by metagenomic analyses. This species includes all bacteria with genomes that show ≥95% average nucleotide identity (ANI) to the type genome for the species to which we have assigned the MAG ID E3_MB2_98 and which is available via NCBI BioSample SAMN18472500. The GC content of the type genome is 54.9 % and the genome length is 1.48 Mbp.

### Description of *Candidatus* Cryptobacteroides caccocaballi sp. nov

*Candidatus* Cryptobacteroides caccocaballi (cac.co.ca.bal’li. Gr. fem. n. *kakke*, faeces; L. masc. n. *caballus*, a horse; N.L. gen. n. *caccocaballi*, associated with the faeces of horses) A bacterial species identified by metagenomic analyses. This species includes all bacteria with genomes that show ≥95% average nucleotide identity (ANI) to the type genome for the species to which we have assigned the MAG ID E4_MB2_58 and which is available via NCBI BioSample SAMN18472527. The GC content of the type genome is 51.8 % and the genome length is 2.22 Mbp.

### Description of *Candidatus* Cryptobacteroides choladohippi sp. nov

*Candidatus* Cryptobacteroides choladohippi (cho.la.do.hip’pi. Gr. fem. n. *kholas*, *kholados*, guts; Gr. masc./fem. n. *hippos*a horse; N.L. gen. n. *choladohippi*, associated with the horse gut)

A bacterial species identified by metagenomic analyses. This species includes all bacteria with genomes that show ≥95% average nucleotide identity (ANI) to the type genome for the species to which we have assigned the MAG ID E1_MB2_55 and which is available via NCBI BioSample SAMN18472464. The GC content of the type genome is 54.2 % and the genome length is 2.24 Mbp.

### Description of *Candidatus* Cryptobacteroides equifaecalis sp. nov

*Candidatus* Cryptobacteroides equifaecalis (e.qui.fae.ca’lis. L. masc. n. *equus*, a horse; N.L. masc. adj. *faecalis*, faecal; N.L. masc. adj. *equifaecalis*, associated with the faeces of horses)

A bacterial species identified by metagenomic analyses. This species includes all bacteria with genomes that show ≥95% average nucleotide identity (ANI) to the type genome for the species to which we have assigned the MAG ID E4_MB2_98 and which is available via NCBI BioSample SAMN18472533. The GC content of the type genome is 52.5 % and the genome length is 1.61 Mbp.

### Description of *Candidatus* Cryptobacteroides faecihippi sp. nov

*Candidatus* Cryptobacteroides faecihippi (fae.ci.hip’pi. L. fem. n. *faex*, *faeces*, *dregs;* Gr. masc./fem. n. *hippos*a horse; N.L. gen. n. *faecihippi*, associated with the faeces of horses) A bacterial species identified by metagenomic analyses. This species includes all bacteria with genomes that show ≥95% average nucleotide identity (ANI) to the type genome for the species to which we have assigned the MAG ID E1_MB2_112 and which is available via NCBI BioSample SAMN18472461. The GC content of the type genome is 54.8 % and the genome length is 2.25 Mbp.

### Description of *Candidatus* Cryptobacteroides fimicaballi sp. nov

*Candidatus* Cryptobacteroides fimicaballi (fi.mi.ca.bal’li. L. masc. n. *fimus*, dung; L. masc. n. *caballus*, a horse; N.L. gen. n. *fimicaballi*, associated with the faeces of horses)

A bacterial species identified by metagenomic analyses. This species includes all bacteria with genomes that show ≥95% average nucleotide identity (ANI) to the type genome for the species to which we have assigned the MAG ID E3_MB2_135 and which is available via NCBI BioSample SAMN18472490. The GC content of the type genome is 51 % and the genome length is 1.33 Mbp.

### Description of *Candidatus* Cryptobacteroides onthequi sp. nov

*Candidatus* Cryptobacteroides onthequi (onth.e’qui. Gr. masc. n. *onthos*, dung; L. masc. n. *equus*, a horse; N.L. gen. n. *onthequi*, associated with the faeces of horses)

A bacterial species identified by metagenomic analyses. This species includes all bacteria with genomes that show ≥95% average nucleotide identity (ANI) to the type genome for the species to which we have assigned the MAG ID E1_MB2_10 and which is available via NCBI BioSample SAMN18472459. The GC content of the type genome is 53.4 % and the genome length is 2.96 Mbp.

### Description of *Candidatus* Cryptobacteroides onthocaballi sp. nov

*Candidatus* Cryptobacteroides onthocaballi (on.tho.ca.bal’li. Gr. masc. n. *onthos*, dung; L. masc. n. *caballus*, a horse; N.L. gen. n. *onthocaballi*, associated with the faeces of horses) A bacterial species identified by metagenomic analyses. This species includes all bacteria with genomes that show ≥95% average nucleotide identity (ANI) to the type genome for the species to which we have assigned the MAG ID E3_MB2_147 and which is available via NCBI BioSample SAMN18472491. The GC content of the type genome is 52 % and the genome length is 1.47 Mbp.

### Description of *Candidatus* Egerieousia equi sp. nov

*Candidatus* Egerieousia equi (e’qui. L. gen. masc. n. *equi,* of a horse)

A bacterial species identified by metagenomic analyses. This species includes all bacteria with genomes that show ≥95% average nucleotide identity (ANI) to the type genome for the species to which we have assigned the MAG ID E4_MB2_106 and which is available via NCBI BioSample SAMN18472513. The GC content of the type genome is 46.4 % and the genome length is 1.92 Mbp.

### Description of *Candidatus* Enterousia merdequi sp. nov

*Candidatus* Enterousia merdequi (merd.e’qui. L. fem. n. *merda*, faeces; L. masc. n. *equus*, a horse; N.L. gen. n. *merdequi*, associated with the faeces of horses)

A bacterial species identified by metagenomic analyses. This species includes all bacteria with genomes that show ≥95% average nucleotide identity (ANI) to the type genome for the species to which we have assigned the MAG ID E3_MB2_90 and which is available via NCBI BioSample SAMN18472497. The GC content of the type genome is 33.9 % and the genome length is 0.74 Mbp.

### Description of *Candidatus* Enterousia onthequi sp. nov

*Candidatus* Enterousia onthequi (onth.e’qui. Gr. masc. n. *onthos*, dung; L. masc. n. *equus*, a horse; N.L. gen. n. *onthequi*, associated with the faeces of horses)

A bacterial species identified by metagenomic analyses. This species includes all bacteria with genomes that show ≥95% average nucleotide identity (ANI) to the type genome for the species to which we have assigned the MAG ID E5_MB2_19 and which is available via NCBI BioSample SAMN18472546. The GC content of the type genome is 38.8 % and the genome length is 0.88 Mbp.

### Description of *Candidatus* Enterousia scatequi sp. nov

*Candidatus* Enterousia scatequi (scat.e’qui. Gr. neut. n. *skor*, *skatos,* dung; L. masc. n. *equus*, a horse; N.L. gen. n. *scatequi*, associated with the faeces of horses)

A bacterial species identified by metagenomic analyses. This species includes all bacteria with genomes that show ≥95% average nucleotide identity (ANI) to the type genome for the species to which we have assigned the MAG ID E5_MB2_120 and which is available via NCBI BioSample SAMN18472543. The GC content of the type genome is 39.9 % and the genome length is 0.76 Mbp.

### Description of *Candidatus* Equadaptatus gen. nov

*Candidatus* Equadaptaus (Equ.a.dap.ta’tus. L. masc. n. *equus*, a horse; L. masc. perf. part. *adaptatus*, adapted to; N.L. masc. n. *Equiadaptatus*, a microbe associated with horses)

A bacterial genus identified by metagenomic analyses. The genus includes all bacteria with genomes that show ≥60% average amino acid identity (AAI) to the genome of the type strain from the type species *Candidatus* Equadaptatus faecalis. This genus has been assigned by GTDB-Tk v1.3.0 working on GTDB Release 05-RS95 (39, 79) to the order *Synergistales* and to the family *Synergistaceae*.

### Description of *Candidatus* Equadaptatus faecalis sp. nov

*Candidatus* Equadaptatus faecalis (fae.ca’lis. N.L. masc. adj. *faecalis*, faecal)

A bacterial species identified by metagenomic analyses. This species includes all bacteria with genomes that show ≥95% average nucleotide identity (ANI) to the type genome for the species to which we have assigned the MAG ID E4_60 and which is available via NCBI BioSample SAMN18472510. The GC content of the type genome is 48.4 % and the genome length is 1.60 Mbp.

### Description of *Candidatus* Equibacterium gen. nov

*Candidatus* Equibacterium (E.qui.bac.te’ri.um. L. masc. n. *equus*, a horse; L. neut. n. *bacterium*, a bacterium; N.L. neut. n. *Equibacterium* a microbe associated with horses)

A bacterial genus identified by metagenomic analyses. The genus includes all bacteria with genomes that show ≥60% average amino acid identity (AAI) to the genome of the type strain from the type species *Candidatus* Equibacterium intestinale. This genus has been assigned by GTDB-Tk v1.3.0 working on GTDB Release 05-RS95 (39, 79) to the order *Bacteroidales* and to the family *UBA932*.

### Description of *Candidatus* Equibacterium intestinale sp. nov

*Candidatus* Equibacterium intestinale (in.tes.ti.na’le.N.L. neut. adj. *intestinale*, pertaining to the intestines)

A bacterial species identified by metagenomic analyses. This species includes all bacteria with genomes that show ≥95% average nucleotide identity (ANI) to the type genome for the species to which we have assigned the MAG ID E5_MB2_82 and which is available via NCBI BioSample SAMN18472552. The GC content of the type genome is 52.3 % and the genome length is 1.76 Mbp.

### Description of *Candidatus* Equicaccousia gen. nov

*Candidatus* Equicaccousia (E.qui.cacc.ou’s.ia. L. masc. n. *equus*, a horse; Gr. fem. n. *kakke*, faeces; Gr. fem. n. *ousia*, an essence; N.L. fem. n. *Equicaccousia*, a microbe associated with horse faeces)

A bacterial genus identified by metagenomic analyses. The genus includes all bacteria with genomes that show ≥60% average amino acid identity (AAI) to the genome of the type strain from the type species *Candidatus* Equicaccousia limihippi. This is a new name for the GTDB alphanumeric genus UMGS1279, which is found in diverse mammalian guts. This genus has been assigned by GTDB-Tk v1.3.0 working on GTDB Release 05-RS95 (39, 79) to the order *Oscillospirales* and to the family *Acutalibacteraceae*.

### Description of *Candidatus* Equicaccousia limihippi sp. nov

*Candidatus* Equicaccousia limihippi (li.mi.hip’pi. L. masc. n. *limus*, dung; Gr. masc./fem. n. *hippos*a horse; N.L. gen. n. *limihippi*, of horse dung)

A bacterial species identified by metagenomic analyses. This species includes all bacteria with genomes that show ≥95% average nucleotide identity (ANI) to the type genome for the species to which we have assigned the MAG ID E2_98 and which is available via NCBI BioSample SAMN18472475. The GC content of the type genome is 44.9 % and the genome length is 1.15 Mbp.

### Description of *Candidatus* Equicola gen. nov

*Candidatus* Equicola (E.qui’co.la. L. masc. n. *equus*, a horse; N.L. masc./fem. suffix *–cola*, an inhabitant; N.L. fem. n. *Equicola*, a microbe associated with horses)

A bacterial genus identified by metagenomic analyses. The genus includes all bacteria with genomes that show ≥60% average amino acid identity (AAI) to the genome of the type strain from the type species *Candidatus* Equicola stercoris. This genus has been assigned by GTDB-Tk v1.3.0 working on GTDB Release 05-RS95 (39, 79) to the order *Bacteroidales* and to the family *Bacteroidaceae*.

### Description of *Candidatus* Equicola faecalis sp. nov

*Candidatus* Equicola faecalis (fae.ca’lis. N.L. fem. adj. *faecalis*, faecal)

A bacterial species identified by metagenomic analyses. This species includes all bacteria with genomes that show ≥95% average nucleotide identity (ANI) to the type genome for the species to which we have assigned the MAG ID E4_176 and which is available via NCBI BioSample SAMN18472505. The GC content of the type genome is 44.8 % and the genome length is 2.09 Mbp.

### Description of *Candidatus* Equicola stercoris sp. nov

*Candidatus* Equicola stercoris (ster’co.ris. L. gen. masc. n. *stercoris*, of dung)

A bacterial species identified by metagenomic analyses. This species includes all bacteria with genomes that show ≥95% average nucleotide identity (ANI) to the type genome for the species to which we have assigned the MAG ID E3_MB2_38 and which is available via NCBI BioSample SAMN18472493. The GC content of the type genome is 42 % and the genome length is 1.75 Mbp.

### Description of *Candidatus* Equihabitans gen. nov

*Candidatus* Equihabitans (E.qui.ha’bi.tans. L. masc. n. *equus*, a horse; L. masc./fem. pres. part. *habitans*, an inhabitant; N.L. fem. n. *Equihabitans*, a microbe associated with horses)

A bacterial genus identified by metagenomic analyses. The genus includes all bacteria with genomes that show ≥60% average amino acid identity (AAI) to the genome of the type strain from the type species *Candidatus* Equihabitans merdae. This genus has been assigned by GTDB-Tk v1.3.0 working on GTDB Release 05-RS95 (39, 79) to the order *Lachnospirales* and to the family *Lachnospiraceae*.

### Description of *Candidatus* Equihabitans merdae sp. nov

*Candidatus* Equihabitans merdae (mer’dae. L. gen. fem. n. *merdae*, of faeces)

A bacterial species identified by metagenomic analyses. This species includes all bacteria with genomes that show ≥95% average nucleotide identity (ANI) to the type genome for the species to which we have assigned the MAG ID E4_98 and which is available via NCBI BioSample SAMN18472512. The GC content of the type genome is 47 % and the genome length is 1.86 Mbp.

### Description of *Candidatus* Equimonas gen. nov

*Candidatus* Equimonas (E.qui.mo’nas. L. masc. n. *equus*, a horse; L. fem. n. *monas*, a monad; N.L. fem. n. *Equimonas*, a microbe associated with horses)

A bacterial genus identified by metagenomic analyses. The genus includes all bacteria with genomes that show ≥60% average amino acid identity (AAI) to the genome of the type strain from the type species *Candidatus* Equimonas enterica. This genus has been assigned by GTDB-Tk v1.3.0 working on GTDB Release 05-RS95 (39, 79) to the order *Bacteroidales* and to the family *Bacteroidaceae*.

### Description of *Candidatus* Equimonas enterica sp. nov

*Candidatus* Equimonas enterica (en.te’ri.ca. Gr. neut. n. *enteron,* gut, bowel, intestine; L.. fem. adj. suff. *-ica,* pertaining to; N.L. fem. adj. *enterica*, pertaining to intestine)

A bacterial species identified by metagenomic analyses. This species includes all bacteria with genomes that show ≥95% average nucleotide identity (ANI) to the type genome for the species to which we have assigned the MAG ID E1_145 and which is available via NCBI BioSample SAMN18472453. The GC content of the type genome is 55.9 % and the genome length is 1.85 Mbp.

### Description of *Candidatus* Equimonas faecalis sp. nov

*Candidatus* Equimonas faecalis (fae.ca’lis. N.L. fem. adj. *faecalis*, faecal)

A bacterial species identified by metagenomic analyses. This species includes all bacteria with genomes that show ≥95% average nucleotide identity (ANI) to the type genome for the species to which we have assigned the MAG ID E1_115 and which is available via NCBI BioSample SAMN18472452. The GC content of the type genome is 55.6 % and the genome length is 2.59 Mbp.

### Description of *Candidatus* Equinaster gen. nov

*Candidatus* Equinaster (E.qui.nas’ter. L. masc. n. *equus*, a horse; Gr. masc. n. naster, an inhabitant; N.L. masc. n. *Equinaster*, a microbe associated with horses)

A bacterial genus identified by metagenomic analyses. The genus includes all bacteria with genomes that show ≥60% average amino acid identity (AAI) to the genome of the type strain from the type species *Candidatus* Equinaster intestinalis. This genus has been assigned by GTDB-Tk v1.3.0 working on GTDB Release 05-RS95 (39, 79) to the order *Oscillospirales* and to the family *Acutalibacteraceae*.

### Description of *Candidatus* Equinaster intestinalis sp. nov

*Candidatus* Equinaster intestinalis (in.tes.ti.na’lis. N.L. masc. adj. *intestinalis*, pertaining to the intestines)

A bacterial species identified by metagenomic analyses. This species includes all bacteria with genomes that show ≥95% average nucleotide identity (ANI) to the type genome for the species to which we have assigned the MAG ID E3_MB2_43 and which is available via NCBI BioSample SAMN18472494. The GC content of the type genome is 43.4 % and the genome length is 1.50 Mbp.

### Description of *Candidatus* Equispira gen. nov

*Candidatus* Equispira (E.qui.spi’ra. L. masc. n. *equus*, a horse; Gr. fem. n. *speira*, a coil, helix; N.L. fem. n. *Equispira*, a helical microbe associated with horses)

A bacterial genus identified by metagenomic analyses. The genus includes all bacteria with genomes that show ≥60% average amino acid identity (AAI) to the genome of the type strain from the type species *Candidatus* Equispira faecalis. This genus has been assigned by GTDB-Tk v1.3.0 working on GTDB Release 05-RS95 (39, 79) to the order *Lachnospirales* and to the family *Lachnospiraceae*.

### Description of *Candidatus* Equispira faecalis sp. nov

*Candidatus* Equispira faecalis (fae.ca’lis. N.L. fem. adj. *faecalis*, faecal)

A bacterial species identified by metagenomic analyses. This species includes all bacteria with genomes that show ≥95% average nucleotide identity (ANI) to the type genome for the species to which we have assigned the MAG ID E5_MB2_109 and which is available via NCBI BioSample SAMN18472542. The GC content of the type genome is 39.2 % and the genome length is 2.33 Mbp.

### Description of *Candidatus* Faecinaster gen. nov

*Candidatus* Faecinaster (Fae.ci.nas’ter. L. fem. n. *faex*, *faecis*, dregs; Gr. masc. n. *naster*, an inhabitant; N.L. masc. n. *Faecinaster*, a microbe associated with faeces)

A bacterial genus identified by metagenomic analyses. The genus includes all bacteria with genomes that show ≥60% average amino acid identity (AAI) to the genome of the type strain from the type species *Candidatus* Faecinaster equi. This is a new name for the GTDB alphanumeric genus UBA6382, which is found in diverse mammalian guts. This genus has been assigned by GTDB-Tk v1.3.0 working on GTDB Release 05-RS95 (39, 79) to the order *Bacteroidales* and to the family *Bacteroidaceae*.

### Description of *Candidatus* Faecinaster equi sp. nov

*Candidatus* Faecinaster equi (e’qui. L. gen. masc. n. *equi*, of a horse)

A bacterial species identified by metagenomic analyses. This species includes all bacteria with genomes that show ≥95% average nucleotide identity (ANI) to the type genome for the species to which we have assigned the MAG ID E3_MB2_9 and which is available via NCBI BioSample SAMN18472496. The GC content of the type genome is 37.2 % and the genome length is 3.36 Mbp.

### Description of *Candidatus* Fiminaster gen. nov

*Candidatus* Fiminaster (Fi.mi.nas’ter. L. neut. n. *fimum*, dung; Gr. masc. n. *naster*, an inhabitant; N.L. masc. n. *Fiminaster*, a microbe associated with faeces)

A bacterial genus identified by metagenomic analyses. The genus includes all bacteria with genomes that show ≥60% average amino acid identity (AAI) to the genome of the type strain from the type species *Candidatus* Fiminaster equi. This is a new name for the GTDB alphanumeric genus UBA3207, which is found in diverse mammalian guts. This genus has been assigned by GTDB-Tk v1.3.0 working on GTDB Release 05-RS95 (39, 79) to the order *RFN20* and to the family *CAG-826*.

### Description of *Candidatus* Fiminaster equi sp. nov

*Candidatus* Fiminaster equi (e’qui. L. gen. masc. n. *equi*, of a horse)

A bacterial species identified by metagenomic analyses. This species includes all bacteria with genomes that show ≥95% average nucleotide identity (ANI) to the type genome for the species to which we have assigned the MAG ID E4_MB2_69 and which is available via NCBI BioSample SAMN18472528. The GC content of the type genome is 34.5 % and the genome length is 0.89 Mbp.

### Description of *Candidatus* Flavobacterium equi sp. nov

*Candidatus* Flavobacterium equi (e’qui. L. gen. masc. n. *equi*, of a horse)

A bacterial species identified by metagenomic analyses. This species includes all bacteria with genomes that show ≥95% average nucleotide identity (ANI) to the type genome for the species to which we have assigned the MAG ID E2_MB2_6 and which is available via NCBI BioSample SAMN18472477. The GC content of the type genome is 37.7 % and the genome length is 2.17 Mbp.

### Description of *Candidatus* Hippenecus gen. nov

*Candidatus* Hippenecus (Hipp.en.e’cus. Gr. masc./fem. n. *hippos*, a horse; N.L. masc. n. *enecus*, an inhabitant; N.L. masc. n. *Hippenecus* a microbe associated with horses)

A bacterial genus identified by metagenomic analyses. The genus includes all bacteria with genomes that show ≥60% average amino acid identity (AAI) to the genome of the type strain from the type species *Candidatus* Hippenecus merdae. This genus has been assigned by GTDB-Tk v1.3.0 working on GTDB Release 05-RS95 (39, 79) to the order *Lachnospirales* and to the family *Lachnospiraceae*.

### Description of *Candidatus* Hippenecus merdae sp. nov

*Candidatus* Hippenecus merdae (mer’dae. L. gen. fem. n. *merdae*, of faeces)

A bacterial species identified by metagenomic analyses. This species includes all bacteria with genomes that show ≥95% average nucleotide identity (ANI) to the type genome for the species to which we have assigned the MAG ID E3_87 and which is available via NCBI BioSample SAMN18472489. The GC content of the type genome is 52.7 % and the genome length is 1.11 Mbp.

### Description of *Candidatus* Hippobium gen. nov

*Candidatus* Hippobium (Hip.po’bi.um. Gr. masc./fem. n. *hippos*, a horse; Gr. masc. n. *bios*, life; N.L. neut. n. *Hippobium*, a microbe associated with horses)

A bacterial genus identified by metagenomic analyses. The genus includes all bacteria with genomes that show ≥60% average amino acid identity (AAI) to the genome of the type strain from the type species *Candidatus* Hippobium faecium. This genus has been assigned by

GTDB-Tk v1.3.0 working on GTDB Release 05-RS95 (39, 79) to the order *UBA5829* and to the family *UBA5829*.

### Description of *Candidatus* Hippobium faecium sp. nov

*Candidatus* Hippobium faecium (fae’ci.um. L. fem. n. *faex*, *dregs*; L. gen. pl. n. *faecium*, of the dregs, of faeces)

A bacterial species identified by metagenomic analyses. This species includes all bacteria with genomes that show ≥95% average nucleotide identity (ANI) to the type genome for the species to which we have assigned the MAG ID E3_206 and which is available via NCBI BioSample SAMN18472485. The GC content of the type genome is 39.1 % and the genome length is 2.12 Mbp.

### Description of *Candidatus* Holdemanella equi sp. nov

*Candidatus* Holdemanella equi (e’qui. L. gen. masc. n. *equi*, of a horse)

A bacterial species identified by metagenomic analyses. This species includes all bacteria with genomes that show ≥95% average nucleotide identity (ANI) to the type genome for the species to which we have assigned the MAG ID E4_26 and which is available via NCBI BioSample SAMN18472508. The GC content of the type genome is 35.6 % and the genome length is 2.44 Mbp.

### Description of *Candidatus* Kurthia equi sp. nov

*Candidatus* Kurthia equi (e’qui. L. gen. masc. n. *equi*, of a horse)

A bacterial species identified by metagenomic analyses. This species includes all bacteria with genomes that show ≥95% average nucleotide identity (ANI) to the type genome for the species to which we have assigned the MAG ID E1_MB2_88 and which is available via NCBI BioSample SAMN18472468. The GC content of the type genome is 35.7 % and the genome length is 3.58 Mbp.

### Description of *Candidatus* Limimonas gen. nov

*Candidatus* Limimonas (Li.mi.mo’nas. L. masc. n. *limus*, dung; L. fem. n. *monas*, a monad; N.L. fem. n. *Limimonas*, a microbe associated with faeces)

A bacterial genus identified by metagenomic analyses. The genus includes all bacteria with genomes that show ≥60% average amino acid identity (AAI) to the genome of the type strain from the type species *Candidatus* Limimonas coprohippi. This is a new name for the GTDB alphanumeric genus UBA1227, which is found in diverse mammalian guts. This genus has been assigned by GTDB-Tk v1.3.0 working on GTDB Release 05-RS95 (39, 79) to the order *Oscillospirales* and to the family *Acutalibacteraceae*.

### Description of *Candidatus* Limimonas coprohippi sp. nov

*Candidatus* Limimonas coprohippi (co.pro.hip’pi. Gr. fem. n. *kopros*, dung; Gr. masc./fem. n. *hippos*, a horse; N.L. gen. n. *coprohippi*, associated with the faeces of horses)

A bacterial species identified by metagenomic analyses. This species includes all bacteria with genomes that show ≥95% average nucleotide identity (ANI) to the type genome for the species to which we have assigned the MAG ID E1_MB2_82 and which is available via NCBI BioSample SAMN18472467. The GC content of the type genome is 40.5 % and the genome length is 1.33 Mbp.

### Description of *Candidatus* Limimonas egerieequi sp. nov

*Candidatus* Limimonas egerieequi (e.ge.ri.e.e’qui. L. fem. n. egeries, *dung;* L. masc. n. *equus*, a horse; N.L. gen. n. *egerieequi*, associated with the faeces of horses)

A bacterial species identified by metagenomic analyses. This species includes all bacteria with genomes that show ≥95% average nucleotide identity (ANI) to the type genome for the species to which we have assigned the MAG ID E5_MB2_129 and which is available via NCBI BioSample SAMN18472544. The GC content of the type genome is 41.8 % and the genome length is 1.70 Mbp.

### Description of *Candidatus* Limimorpha caballi sp. nov

*Candidatus* Limimorpha caballi (ca.bal’li. L. gen. masc. n. *caballi*, of a horse)

A bacterial species identified by metagenomic analyses. This species includes all bacteria with genomes that show ≥95% average nucleotide identity (ANI) to the type genome for the species to which we have assigned the MAG ID E5_119 and which is available via NCBI BioSample SAMN18472534. The GC content of the type genome is 48.3 % and the genome length is 2.76 Mbp.

### Description of *Candidatus* Limimorpha equi sp. nov

*Candidatus* Limimorpha equi (e’qui. L. gen. masc. n. *equi*, of a horse)

A bacterial species identified by metagenomic analyses. This species includes all bacteria with genomes that show ≥95% average nucleotide identity (ANI) to the type genome for the species to which we have assigned the MAG ID E1_MB2_99 and which is available via NCBI BioSample SAMN18472470. The GC content of the type genome is 45.1 % and the genome length is 2.72 Mbp.

### Description of *Candidatus* Liminaster gen. nov

*Candidatus* Liminaster (Li.mi.nas’ter. L. masc. n. *limus*, dung; Gr. masc. n. *naster*, an inhabitant; N.L. masc. n. *Liminaster*, a microbe associated with faeces)

A bacterial genus identified by metagenomic analyses. The genus includes all bacteria with genomes that show ≥60% average amino acid identity (AAI) to the genome of the type strain from the type species *Candidatus* Liminaster caballi. This is a new name for the GTDB alphanumeric genus UBA3663, which is found in diverse mammalian guts. This genus has been assigned by GTDB-Tk v1.3.0 working on GTDB Release 05-RS95 (39, 79) to the order *Bacteroidales* and to the family *UBA3663*.

### Description of *Candidatus* Liminaster caballi sp. nov

*Candidatus* Liminaster caballi (ca.bal’li. L. gen. masc. n. *caballi*, of a horse)

A bacterial species identified by metagenomic analyses. This species includes all bacteria with genomes that show ≥95% average nucleotide identity (ANI) to the type genome for the species to which we have assigned the MAG ID E4_95 and which is available via NCBI BioSample SAMN18472511. The GC content of the type genome is 50.1 % and the genome length is 2.94 Mbp.

### Description of *Candidatus* Merdinaster gen. nov

*Candidatus* Merdinaster (Mer.di.nas’ter. L. fem. n. *merda*, dung; Gr. masc. n. *naster*, an inhabitant; N.L. masc. n. *Merdinaster*, a microbe associated with faeces)

A bacterial genus identified by metagenomic analyses. The genus includes all bacteria with genomes that show ≥60% average amino acid identity (AAI) to the genome of the type strain from the type species *Candidatus* Merdinaster equi. This is a new name for the GTDB alphanumeric genus UBA7050, which is found in diverse mammalian guts. This genus has been assigned by GTDB-Tk v1.3.0 working on GTDB Release 05-RS95 (39, 79) to the order *Lachnospirales* and to the family *Lachnospiraceae*,

### Description of *Candidatus* Merdinaster equi sp. nov

*Candidatus* Merdinaster equi (e’qui. L. gen. masc. n. *equi*, of a horse)

A bacterial species identified by metagenomic analyses. This species includes all bacteria with genomes that show ≥95% average nucleotide identity (ANI) to the type genome for the species to which we have assigned the MAG ID E4_MB2_128 and which is available via NCBI BioSample SAMN18472516. The GC content of the type genome is 40.7 % and the genome length is 1.95 Mbp.

### Description of *Candidatus* Methanocorpusculum equi sp. nov

*Candidatus* Methanocorpusculum equi (e’qui. L. gen. masc. n. *equi*, of a horse)

A bacterial species identified by metagenomic analyses. This species includes all bacteria with genomes that show ≥95% average nucleotide identity (ANI) to the type genome for the species to which we have assigned the MAG ID E2_MB2_79 and which is available via NCBI BioSample SAMN18472479. The GC content of the type genome is 50.2 % and the genome length is 1.15 Mbp.

### Description of *Candidatus* Minthenecus gen. nov

*Candidatus* Minthenecus (Minth.en.e’cus. Gr. masc. n. *minthos*, dung; N.L. masc. n. *enecus*, an inhabitant; N.L. masc. n. *Minthenecus*, a microbe associated with faeces)

A bacterial genus identified by metagenomic analyses. The genus includes all bacteria with genomes that show ≥60% average amino acid identity (AAI) to the genome of the type strain from the type species *Candidatus* Minthenecus merdequi. This is a new name for the GTDB alphanumeric genus SFVR01, which is found in diverse mammalian guts. This genus has been assigned by GTDB-Tk v1.3.0 working on GTDB Release 05-RS95 (39, 79) to the order *Bacteroidales* and to the family *Paludibacteraceae*.

### Description of *Candidatus* Minthenecus merdequi sp. nov

*Candidatus* Minthenecus merdequi (merd.e’qui. L. fem. n. *merda*, faeces; L. masc. n. *equus*, a horse; N.L. gen. n. *merdequi*, associated with the faeces of horses)

A bacterial species identified by metagenomic analyses. This species includes all bacteria with genomes that show ≥95% average nucleotide identity (ANI) to the type genome for the species to which we have assigned the MAG ID E5_MB2_18 and which is available via NCBI BioSample SAMN18472545. The GC content of the type genome is 42.5 % and the genome length is 1.80 Mbp.

### Description of *Candidatus* Minthocola gen. nov

*Candidatus* Minthocola (Min.tho’co.la. Gr. masc. n. *minthos*, dung; N.L. masc./fem. suffix *- cola*, an inhabitant; N.L. fem. n. *Minthocola*, a microbe associated with faeces)

A bacterial genus identified by metagenomic analyses. The genus includes all bacteria with genomes that show ≥60% average amino acid identity (AAI) to the genome of the type strain from the type species *Candidatus* Minthocola equi. This is a new name for the GTDB alphanumeric genus UBA3774, which is found in diverse mammalian guts. This genus has been assigned by GTDB-Tk v1.3.0 working on GTDB Release 05-RS95 (39, 79) to the order *Lachnospirales* and to the family *Lachnospiraceae*.

### Description of *Candidatus* Minthocola equi sp. nov

*Candidatus* Minthocola equi (e’qui. L. gen. masc. n. *equi*, of a horse)

A bacterial species identified by metagenomic analyses. This species includes all bacteria with genomes that show ≥95% average nucleotide identity (ANI) to the type genome for the species to which we have assigned the MAG ID E5_MB2_38 and which is available via NCBI BioSample SAMN18472548. The GC content of the type genome is 45.2 % and the genome length is 1.20 Mbp.

### Description of *Candidatus* Minthomonas gen. nov

*Candidatus* Minthomonas (Min.tho.mo’nas. Gr. masc. n. *minthos*, dung; L. fem. n. *monas*, a monad; N.L. fem. n. *Minthomonas*, a microbe associated with faeces)

A bacterial genus identified by metagenomic analyses. The genus includes all bacteria with genomes that show ≥60% average amino acid identity (AAI) to the genome of the type strain from the type species *Candidatus* Minthomonas equi. This is a new name for the GTDB alphanumeric genus CAG-831, which is found in diverse mammalian guts. This genus has been assigned by GTDB-Tk v1.3.0 working on GTDB Release 05-RS95 (39, 79) to the order *Bacteroidales* and to the family *UBA932*.

### Description of *Candidatus* Minthomonas equi sp. nov

*Candidatus* Minthomonas equi (e’qui. L. gen. masc. n. *equi*, of a horse)

A bacterial species identified by metagenomic analyses. This species includes all bacteria with genomes that show ≥95% average nucleotide identity (ANI) to the type genome for the species to which we have assigned the MAG ID E5_18 and which is available via NCBI

BioSample SAMN18472536. The GC content of the type genome is 47.6 % and the genome length is 1.36 Mbp.

### Description of *Candidatus* Minthosoma gen. nov

*Candidatus* Minthosoma (Min.tho.so’ma. Gr. masc. n. *minthos*, dung; Gr. neut. n. *soma*, a body; N.L. neut. n. *Minthosoma*, a microbe associated with faeces)

A bacterial genus identified by metagenomic analyses. The genus includes all bacteria with genomes that show ≥60% average amino acid identity (AAI) to the genome of the type strain from the type species *Candidatus* Minthosoma caballi. *This* is a new name for the GTDB alphanumeric genus UBA4334, which is found in diverse mammalian guts. This genus has been assigned by GTDB-Tk v1.3.0 working on GTDB Release 05-RS95 (39, 79) to the order *Bacteroidales* and to the family *Bacteroidaceae*

### Description of *Candidatus* Minthosoma caballi sp. nov

*Candidatus* Minthosoma caballi (ca.bal’li. L. gen. masc. n. *caballi*, of a horse)

A bacterial species identified by metagenomic analyses. This species includes all bacteria with genomes that show ≥95% average nucleotide identity (ANI) to the type genome for the species to which we have assigned the MAG ID E5_9 and which is available via NCBI BioSample SAMN18472539. The GC content of the type genome is 44.2 % and the genome length is 3.21 Mbp.

### Description of *Candidatus* Minthosoma equi sp. nov

*Candidatus* Minthosoma equi (e’qui. L. gen. masc. n. *equi*, of a horse)

A bacterial species identified by metagenomic analyses. This species includes all bacteria with genomes that show ≥95% average nucleotide identity (ANI) to the type genome for the species to which we have assigned the MAG ID E4_MB2_18 and which is available via NCBI BioSample SAMN18472519. The GC content of the type genome is 44.1 % and the genome length is 3.51 Mbp.

### Description of *Candidatus* Minthousia gen. nov

*Candidatus* Minthousia (Minth.ou’s.ia. Gr. masc. n. *minthos*, dung; Gr. fem. n. *ousia*, an essence; N.L. fem. n. *Minthousia*, a microbe associated with faeces)

A bacterial genus identified by metagenomic analyses. The genus includes all bacteria with genomes that show ≥60% average amino acid identity (AAI) to the genome of the type strain from the type species *Candidatus* Minthousia equi. This is a new name for the GTDB alphanumeric genus UBA4293, which is found in diverse mammalian guts. This genus has been assigned by GTDB-Tk v1.3.0 working on GTDB Release 05-RS95 (39, 79) to the order *Bacteroidales* and to the family *Bacteroidaceae*.

### Description of *Candidatus* Minthousia equi sp. nov

*Candidatus* Minthousia equi (e’qui. L. gen. masc. n. *equi*, of a horse)

A bacterial species identified by metagenomic analyses. This species includes all bacteria with genomes that show ≥95% average nucleotide identity (ANI) to the type genome for the species to which we have assigned the MAG ID E4_55 and which is available via NCBI BioSample SAMN18472509. The GC content of the type genome is 42.9 % and the genome length is 2.61 Mbp.

### Description of *Candidatus* Mogibacterium equifaecale sp. nov

*Candidatus* Mogibacterium equifaecale (e.qui.fae.ca’le. L. masc. n. *equus*, a horse; N.L. neut. adj. *faecale*, faecal N.L. neut. adj. *equifaecale*, associated with the faeces of horses) A bacterial species identified by metagenomic analyses. This species includes all bacteria with genomes that show ≥95% average nucleotide identity (ANI) to the type genome for the species to which we have assigned the MAG ID E4_MB2_51 and which is available via NCBI BioSample SAMN18472526. The GC content of the type genome is 43.4 % and the genome length is 1.39 Mbp.

### Description of *Candidatus* Mogibacterium onthequi sp. nov

*Candidatus* Mogibacterium onthequi (onth.e’qui. Gr. masc. n. *onthos*, dung; L. masc. n. *equus*, a horse; N.L. gen. n. *onthequi*, associated with the faeces of horses)

A bacterial species identified by metagenomic analyses. This species includes all bacteria with genomes that show ≥95% average nucleotide identity (ANI) to the type genome for the species to which we have assigned the MAG ID E4_MB2_84 and which is available via NCBI BioSample SAMN18472530. The GC content of the type genome is 43.8 % and the genome length is 1.61 Mbp.

### Description of *Candidatus* Mogibacterium scatequi sp. nov

*Candidatus* Mogibacterium scatequi (scat.e’qui. Gr. neut. n. *skor*, *skatos,* dung; L. masc. n. *equus*, a horse; N.L. gen. n. *scatequi*, associated with the faeces of horses)

A bacterial species identified by metagenomic analyses. This species includes all bacteria with genomes that show ≥95% average nucleotide identity (ANI) to the type genome for the species to which we have assigned the MAG ID E4_MB2_90 and which is available via NCBI BioSample SAMN18472532. The GC content of the type genome is 45.2 % and the genome length is 1.45 Mbp.

### Description of *Candidatus* Mycoplasma equi sp. nov

*Candidatus* Mycoplasma equi (e’qui. L. gen. masc. n. *equi*, of a horse)

A bacterial species identified by metagenomic analyses. This species includes all bacteria with genomes that show ≥95% average nucleotide identity (ANI) to the type genome for the species to which we have assigned the MAG ID E4_MB2_29 and which is available via NCBI BioSample SAMN18472521. The GC content of the type genome is 31.4 % and the genome length is 0.64 Mbp.

### Description of *Candidatus* Onthonaster gen. nov

*Candidatus* Onthonaster (On.tho.nas’ter. Gr. masc. n. *onthos*, dung; Gr. masc. n. *naster*, an inhabitant; N.L. masc. n. *Onthonaster*, a microbe associated with faeces)

A bacterial genus identified by metagenomic analyses. The genus includes all bacteria with genomes that show ≥60% average amino acid identity (AAI) to the genome of the type strain from the type species *Candidatus* Onthonaster equi. *This* is a new name for the GTDB alphanumeric genus YIM-102668, which is found in diverse mammalian guts. This genus has been assigned by GTDB-Tk v1.3.0 working on GTDB Release 05-RS95 (39, 79) to the order *Flavobacteriales* and to the family *Weeksellaceae*

### Description of *Candidatus* Onthonaster equi sp. nov

*Candidatus* Onthonaster equi (e’qui. L. gen. masc. n. *equi*, of a horse)

A bacterial species identified by metagenomic analyses. This species includes all bacteria with genomes that show ≥95% average nucleotide identity (ANI) to the type genome for the species to which we have assigned the MAG ID E1_98 and which is available via NCBI BioSample SAMN18472458. This is a new name for the alphanumeric GTDB species sp003687725, which is found in diverse mammalian guts. The GC content of the type genome is 31.1 % and the genome length is 2.30 Mbp.

### Description of *Candidatus* Phascolarctobacterium caballi sp. nov

*Candidatus* Phascolarctobacterium caballi (ca.bal’li. L. gen. masc. n. *caballi*, of a horse)

A bacterial species identified by metagenomic analyses. This species includes all bacteria with genomes that show ≥95% average nucleotide identity (ANI) to the type genome for the species to which we have assigned the MAG ID E4_135 and which is available via NCBI BioSample SAMN18472503. The GC content of the type genome is 39.4 % and the genome length is 1.56 Mbp.

### Description of *Candidatus* Phascolarctobacterium equi sp. nov

*Candidatus* Phascolarctobacterium equi (e’qui. L. gen. masc. n. *equi*, of a horse)

A bacterial species identified by metagenomic analyses. This species includes all bacteria with genomes that show ≥95% average nucleotide identity (ANI) to the type genome for the species to which we have assigned the MAG ID E2_44 and which is available via NCBI BioSample SAMN18472473. The GC content of the type genome is 46.7 % and the genome length is 0.93 Mbp.

### Description of *Candidatus* Physcocola gen. nov

*Candidatus* Physcocola (Phys.co’co.la. Gr. fem. n. *physke*, the colon; N.L. masc./fem. Suffix *–cola*, an inhabitant; N.L. fem. n. *Physcocola*, a microbe associated with the large intestine) A bacterial genus identified by metagenomic analyses. The genus includes all bacteria with genomes that show ≥60% average amino acid identity (AAI) to the genome of the type strain from the type species *Candidatus* Physcocola equi. This is a new name for the GTDB alphanumeric genus UBA4345, which is found in diverse mammalian guts. This genus has been assigned by GTDB-Tk v1.3.0 working on GTDB Release 05-RS95 (39, 79) to the order *Bacteroidales* and to the family *Paludibacteraceae*.

### Description of *Candidatus* Physcocola equi sp. nov

*Candidatus* Physcocola equi (e’qui. L. gen. masc. n. *equi*, of a horse)

A bacterial species identified by metagenomic analyses. This species includes all bacteria with genomes that show ≥95% average nucleotide identity (ANI) to the type genome for the species to which we have assigned the MAG ID E4_MB2_42 and which is available via NCBI BioSample SAMN18472523. The GC content of the type genome is 43.3 % and the genome length is 2.99 Mbp.

### Description of *Candidatus* Physcosoma gen. nov

*Candidatus* Physcosoma (Phys.co.so’ma. Gr. fem. n. *physke*, the colon; Gr. neut. n. *soma*, a body; N.L. neut. n. *Physcosoma*, a microbe associated with the large intestine)

A bacterial genus identified by metagenomic analyses. The genus includes all bacteria with genomes that show ≥60% average amino acid identity (AAI) to the genome of the type strain from the type species *Candidatus* Physcosoma equi. This is a new name for the GTDB alphanumeric genus UBA5920, which is found in diverse mammalian guts. This genus has been assigned by GTDB-Tk v1.3.0 working on GTDB Release 05-RS95 (39, 79) to the order *Sphaerochaetales* and to the family *Sphaerochaetaceae*.

### Description of *Candidatus* Physcosoma equi sp. nov

*Candidatus* Physcosoma equi (e’qui. L. gen. masc. n. *equi*, of a horse)

A bacterial species identified by metagenomic analyses. This species includes all bacteria with genomes that show ≥95% average nucleotide identity (ANI) to the type genome for the species to which we have assigned the MAG ID E4_160 and which is available via NCBI BioSample SAMN18472504. The GC content of the type genome is 49.1 % and the genome length is 2.06 Mbp.

### Description of *Candidatus* Physcousia gen. nov

*Candidatus* Physcousia (Physc.ou’si.a. Gr. fem. n. *physke* the colon; Gr. fem. n. *ousia,* an essence.e; N.L. fem. n. *Physcousia*, a microbe associated with the large intestine)

A bacterial genus identified by metagenomic analyses. The genus includes all bacteria with genomes that show ≥60% average amino acid identity (AAI) to the genome of the type strain from the type species *Candidatus* Physcousia caballi. This is a new name for the GTDB alphanumeric genus UBA4372, which is found in diverse mammalian guts. This genus has been assigned by GTDB-Tk v1.3.0 working on GTDB Release 05-RS95 (39, 79) to the order *Bacteroidales* and to the family *Bacteroidaceae*.

### Description of *Candidatus* Physcousia caballi sp. nov

*Candidatus* Physcousia caballi (ca.bal’li. L. gen. masc. n. *caballi*, of a horse)

A bacterial species identified by metagenomic analyses. This species includes all bacteria with genomes that show ≥95% average nucleotide identity (ANI) to the type genome for the species to which we have assigned the MAG ID E4_MB2_73 and which is available via NCBI BioSample SAMN18472529. The GC content of the type genome is 50.5 % and the genome length is 3.81 Mbp.

### Description of *Candidatus* Physcousia equi sp. nov

*Candidatus* Physcousia equi (e’qui. L. gen. masc. n. *equi*, of a horse)

A bacterial species identified by metagenomic analyses. This species includes all bacteria with genomes that show ≥95% average nucleotide identity (ANI) to the type genome for the species to which we have assigned the MAG ID E4_MB2_112 and which is available via NCBI BioSample SAMN18472514. The GC content of the type genome is 52.4 % and the genome length is 2.43 Mbp.

### Description of *Candidatus* Prevotella equi sp. nov

*Candidatus* Prevotella equi (e’qui. L. gen. masc. n. *equi*, of a horse)

A bacterial species identified by metagenomic analyses. This species includes all bacteria with genomes that show ≥95% average nucleotide identity (ANI) to the type genome for the species to which we have assigned the MAG ID E4_23 and which is available via NCBI BioSample SAMN18472507. The GC content of the type genome is 44.5 % and the genome length is 3.45 Mbp.

### Description of *Candidatus* Ruminococcus equi sp. nov

*Candidatus* Ruminococcus equi (e’qui. L. gen. masc. n. *equi*, of a horse)

A bacterial species identified by metagenomic analyses. This species includes all bacteria with genomes that show ≥95% average nucleotide identity (ANI) to the type genome for the species to which we have assigned the MAG ID E3_41 and which is available via NCBI BioSample SAMN18472487. GTDB has assigned. This species to a genus marked with an alphabetical suffix. However, as this genus designation cannot be incorporated into a well- formed binomial, in naming. This species, we have used the current validly published name for the genus. The GC content of the type genome is 39.9 % and the genome length is 1.73 Mbp.

### Description of *Candidatus* Scatohabitans gen. nov

*Candidatus* Scatohabitans (Sca.to.ha’bi.tans. Gr. neut. n. *skor*, *skatos*, dung; L. masc./fem. pres. part. *habitans*, an inhabitant; N.L. fem. n. *Scatohabitans* a microbe associated with faeces)

A bacterial genus identified by metagenomic analyses. The genus includes all bacteria with genomes that show ≥60% average amino acid identity (AAI) to the genome of the type strain from the type species *Candidatus* Scatohabitans aphodohippi. This is a new name for the GTDB alphanumeric genus C941, which is found in diverse mammalian guts. This genus has been assigned by GTDB-Tk v1.3.0 working on GTDB Release 05-RS95 (39, 79) to the order *Bacteroidales* and to the family *Muribaculaceae*.

### Description of *Candidatus* Scatohabitans aphodohippi sp. nov

*Candidatus* Scatohabitans aphodohippi (aph.o.do.hip’pi. Gr. fem. n. *aphodos*, dung; Gr. masc./fem. n. *hippos*, a horse; N.L. gen. n. *aphodohippi*, associated with the faeces of horses)

A bacterial species identified by metagenomic analyses. This species includes all bacteria with genomes that show ≥95% average nucleotide identity (ANI) to the type genome for the species to which we have assigned the MAG ID E3_0 and which is available via NCBI BioSample SAMN18472480. The GC content of the type genome is 50 % and the genome length is 2.49 Mbp.

### Description of *Candidatus* Scatohabitans fimicaballi sp. nov

*Candidatus* Scatohabitans fimicaballi (fi.mi.ca.bal’li. L. masc. n. *fimus*, dung; L. masc. n. *caballus*, a horse; N.L. gen. n. *fimicaballi*, associated with the faeces of horses)

A bacterial species identified by metagenomic analyses. This species includes all bacteria with genomes that show ≥95% average nucleotide identity (ANI) to the type genome for the species to which we have assigned the MAG ID E4_193 and which is available via NCBI BioSample SAMN18472506. The GC content of the type genome is 48.1 % and the genome length is 2.46 Mbp.

### Description of *Candidatus* Scatohabitans limicaballi sp. nov

*Candidatus* Scatohabitans limicaballi (li.mi.ca.bal’li. L. masc. n. *limus*, dung; L. masc. n. *caballus*, a horse; N.L. gen. n. *limicaballi*, associated with the faeces of horses)

A bacterial species identified by metagenomic analyses. This species includes all bacteria with genomes that show ≥95% average nucleotide identity (ANI) to the type genome for the species to which we have assigned the MAG ID E2_8 and which is available via NCBI BioSample SAMN18472474. The GC content of the type genome is 50.4 % and the genome length is 3.16 Mbp.

### Description of *Candidatus* Scatonaster gen. nov

*Candidatus* Scatonaster (Sca.to.nas’ter. Gr. neut. n. *skor*, *skatos*, dung; Gr. masc. n. *naster*, an inhabitant; N.L. masc. n. *Scatonaster* a microbe associated with faeces)

A bacterial genus identified by metagenomic analyses. The genus includes all bacteria with genomes that show ≥60% average amino acid identity (AAI) to the genome of the type strain from the type species *Candidatus* Scatonaster coprocaballi. This is a new name for the GTDB alphanumeric genus Firm-16, which is found in diverse mammalian guts. This genus has been assigned by GTDB-Tk v1.3.0 working on GTDB Release 05-RS95 (39, 79) to the order *Saccharofermentanales* and to the family *Saccharofermentanaceae*.

### Description of *Candidatus* Scatonaster coprocaballi sp. nov

*Candidatus* Scatonaster coprocaballi (co.pro.ca.bal’li. Gr. fem. n. *kopros*, dung; L. masc. n. *caballus*, a horse; N.L. gen. n. *coprocaballi*, associated with the faeces of horses)

A bacterial species identified by metagenomic analyses. This species includes all bacteria with genomes that show ≥95% average nucleotide identity (ANI) to the type genome for the species to which we have assigned the MAG ID E5_MB2_10 and which is available via NCBI BioSample SAMN18472540. The GC content of the type genome is 46.9 % and the genome length is 2.23 Mbp.

### Description of *Candidatus* Scybalocola gen. nov

*Candidatus* Scybalocola (Scy.ba.lo’co.la. Gr. neut. n. *skybalon*, dung; N.L. masc./fem. Suffix *–cola*, an inhabitant; N.L. fem. n. *Scybalocola* a microbe associated with faeces)

A bacterial genus identified by metagenomic analyses. The genus includes all bacteria with genomes that show ≥60% average amino acid identity (AAI) to the genome of the type strain from the type species *Candidatus* Scybalocola fimicaballi. This is a new name for the GTDB alphanumeric genus UBA1723, which is found in diverse mammalian guts. This genus has been assigned by GTDB-Tk v1.3.0 working on GTDB Release 05-RS95 (39, 79) to the order *Bacteroidales* and to the family *Paludibacteraceae*.

### Description of *Candidatus* Scybalocola fimicaballi sp. nov

*Candidatus* Scybalocola fimicaballi (fi.mi.ca.bal’li. L. masc. n. *fimus*, dung; L. masc. n. *caballus*, a horse; N.L. gen. n. *fimicaballi*, associated with the faeces of horses)

A bacterial species identified by metagenomic analyses. This species includes all bacteria with genomes that show ≥95% average nucleotide identity (ANI) to the type genome for the species to which we have assigned the MAG ID E1_25 and which is available via NCBI BioSample SAMN18472456. This is a new name for the alphanumeric GTDB species sp002317115, which is found in diverse mammalian guts. The GC content of the type genome is 41.7 % and the genome length is 3.11 Mbp.

### Description of *Candidatus* Scybalousia gen. nov

*Candidatus* Scybalousia (Scy.bal.ou’s.ia. Gr. neut. n. *skybalon*, dung; Gr. fem. n. *ousia*, an essence; N.L. fem n. *Scybalousia*, a microbe associated with faeces)

A bacterial genus identified by metagenomic analyses. The genus includes all bacteria with genomes that show ≥60% average amino acid identity (AAI) to the genome of the type strain from the type species *Candidatus* Scybalousia scubalohippi. This is a new name for the GTDB alphanumeric genus Phil12, which is found in diverse mammalian guts. This genus has been assigned by GTDB-Tk v1.3.0 working on GTDB Release 05-RS95 (39, 79) to the order *Bacteroidales* and to the family *P3*.

### Description of *Candidatus* Scybalousia scybalohippi sp. nov

*Candidatus* Scybalousia scybalohippi (scy.ba.lo.hip’pi. Gr. neut. n. *skybalon*, dung; Gr. masc./fem. n. *hippos*, a horse; N.L. gen. n. *scybalohippi*, associated with the faeces of horses)

A bacterial species identified by metagenomic analyses. This species includes all bacteria with genomes that show ≥95% average nucleotide identity (ANI) to the type genome for the species to which we have assigned the MAG ID E3_144 and which is available via NCBI BioSample SAMN18472482. The GC content of the type genome is 35.4 % and the genome length is 2.63 Mbp.

### Description of *Candidatus* Stomatobaculum equi sp. nov

*Candidatus* Stomatobaculum equi (e’qui. L. gen. masc. n. *equi*, of a horse)

A bacterial species identified by metagenomic analyses. This species includes all bacteria with genomes that show ≥95% average nucleotide identity (ANI) to the type genome for the species to which we have assigned the MAG ID E1_MB2_36 and which is available via NCBI BioSample SAMN18472462. GTDB has assigned. This species to a genus marked with an alphabetical suffix. However, as this genus designation cannot be incorporated into a well-formed binomial, in naming. This species, we have used the current validly published name for the genus. The GC content of the type genome is 44.2 % and the genome length is 1.44 Mbp.

### Description of *Candidatus* Treponema caballi sp. nov

*Candidatus* Treponema caballi (ca.bal’li. L. gen. masc. n. *caballi*, of a horse)

A bacterial species identified by metagenomic analyses. This species includes all bacteria with genomes that show ≥95% average nucleotide identity (ANI) to the type genome for the species to which we have assigned the MAG ID E1_106 and which is available via NCBI BioSample SAMN18472451. GTDB has assigned. This species to a genus marked with an alphabetical suffix. However, as this genus designation cannot be incorporated into a well- formed binomial, in naming. This species, we have used the current validly published name for the genus. The GC content of the type genome is 47.1 % and the genome length is 2.91 Mbp.

### Description of *Candidatus* Treponema equi sp. nov

*Candidatus* Treponema equi (e’qui. L. gen. masc. n. *equi*, of a horse)

A bacterial species identified by metagenomic analyses. This species includes all bacteria with genomes that show ≥95% average nucleotide identity (ANI) to the type genome for the species to which we have assigned the MAG ID E4_MB2_46 and which is available via NCBI BioSample SAMN18472525. GTDB has assigned. This species to a genus marked with an alphabetical suffix. However, as this genus designation cannot be incorporated into a well-formed binomial, in naming. This species, we have used the current validly published name for the genus. The GC content of the type genome is 44.3 % and the genome length is 1.79 Mbp.

### Description of *Candidatus* Treponema equifaecale sp. nov

*Candidatus* Treponema equifaecale (e.qui.fae.ca’le. L. masc. n. *equus*, a horse; N.L. neut. adj. *faecale*, faecal; N.L. neut. adj *.equifaecale*, associated with the faeces of horses)

A bacterial species identified by metagenomic analyses. This species includes all bacteria with genomes that show ≥95% average nucleotide identity (ANI) to the type genome for the species to which we have assigned the MAG ID E4_MB2_2 and which is available via NCBI BioSample SAMN18472520. GTDB has assigned. This species to a genus marked with an alphabetical suffix. However, as this genus designation cannot be incorporated into a well- formed binomial, in naming. This species, we have used the current validly published name for the genus. The GC content of the type genome is 40.2 % and the genome length is 2.81 Mbp.

### Description of *Candidatus* Treponema merdequi sp. nov

*Candidatus* Treponema merdequi (merd.e’qui. L. fem. n. *merda*, faeces; L. masc. n. *equus*, a horse; N.L. gen. n. *merdequi*, associated with the faeces of horses)

A bacterial species identified by metagenomic analyses. This species includes all bacteria with genomes that show ≥95% average nucleotide identity (ANI) to the type genome for the species to which we have assigned the MAG ID E5_50 and which is available via NCBI BioSample SAMN18472538. GTDB has assigned. This species to a genus marked with an alphabetical suffix. However, as this genus designation cannot be incorporated into a well- formed binomial, in naming. This species, we have used the current validly published name for the genus. The GC content of the type genome is 35.8 % and the genome length is 2.70 Mbp.

### Description of *Candidatus* Treponema scatequi sp. nov

*Candidatus* Treponema scatequi (scat.e’qui. Gr. neut. n. *skor*, *skatos*, dung; L. masc. n. *equus*, a horse; N.L. gen. n. *scatequi*, associated with the faeces of horses)

A bacterial species identified by metagenomic analyses. This species includes all bacteria with genomes that show ≥95% average nucleotide identity (ANI) to the type genome for the species to which we have assigned the MAG ID E1_MB2_111 and which is available via NCBI BioSample SAMN18472460. GTDB has assigned. This species to a genus marked with an alphabetical suffix. However, as this genus designation cannot be incorporated into a well-formed binomial, in naming this species, we have used the current validly published name for the genus. The GC content of the type genome is 38.4 % and the genome length is

2.31 Mbp.

### A newly named class within the *Armatimonadetes*

One of our MAGs—and the associated metagenomic species, which we have called *Ca.* Hippobium faecium—was assigned to an unnamed family within the recently named phylum *Armatimonadetes* (also called *Armatimonadota;* previously known as OP10) (80). Scrutiny of the NCBI database in March 2021 reveals that no genome assemblies and only twenty-five 16S gene sequences linked to this phylum originate from the vertebrate gut, which include sequences from cattle (81), pigs (82–84), catfish (85) and migratory passerines.

Phylogenetic analysis placed the new species on a deep branch within the *Armatimonadetes*, clustering with a solitary bioreactor-derived metagenome-assembled genome UBA5829 (GCA_002431715.1) assigned by GTDB to its own class, order, family and genus, all given the same alphanumeric designation UBA5829. Sequence comparisons between the genomes of *Ca.* H. faecium UBA5829 report an AAI of 51% (S7 Table), which suggests that they sit within the same class, which we propose should be named *Ca*.

Hippobiia.

### Distribution and metabolism

Our de-replicated high- and medium-quality MAGs account for 18% (±5%) of our host-depleted metagenomic reads. Distribution analysis identified 17 species present at ≥ 1x coverage in all samples, spanning four bacterial phyla and the archaea (Fig 2a and S8 Table). No species were present at ≥10x coverage in all samples. Species quantification shows a steady incline in the cumulative number of species identified with each consecutive sample (Fig 2b).

**Fig 2.**
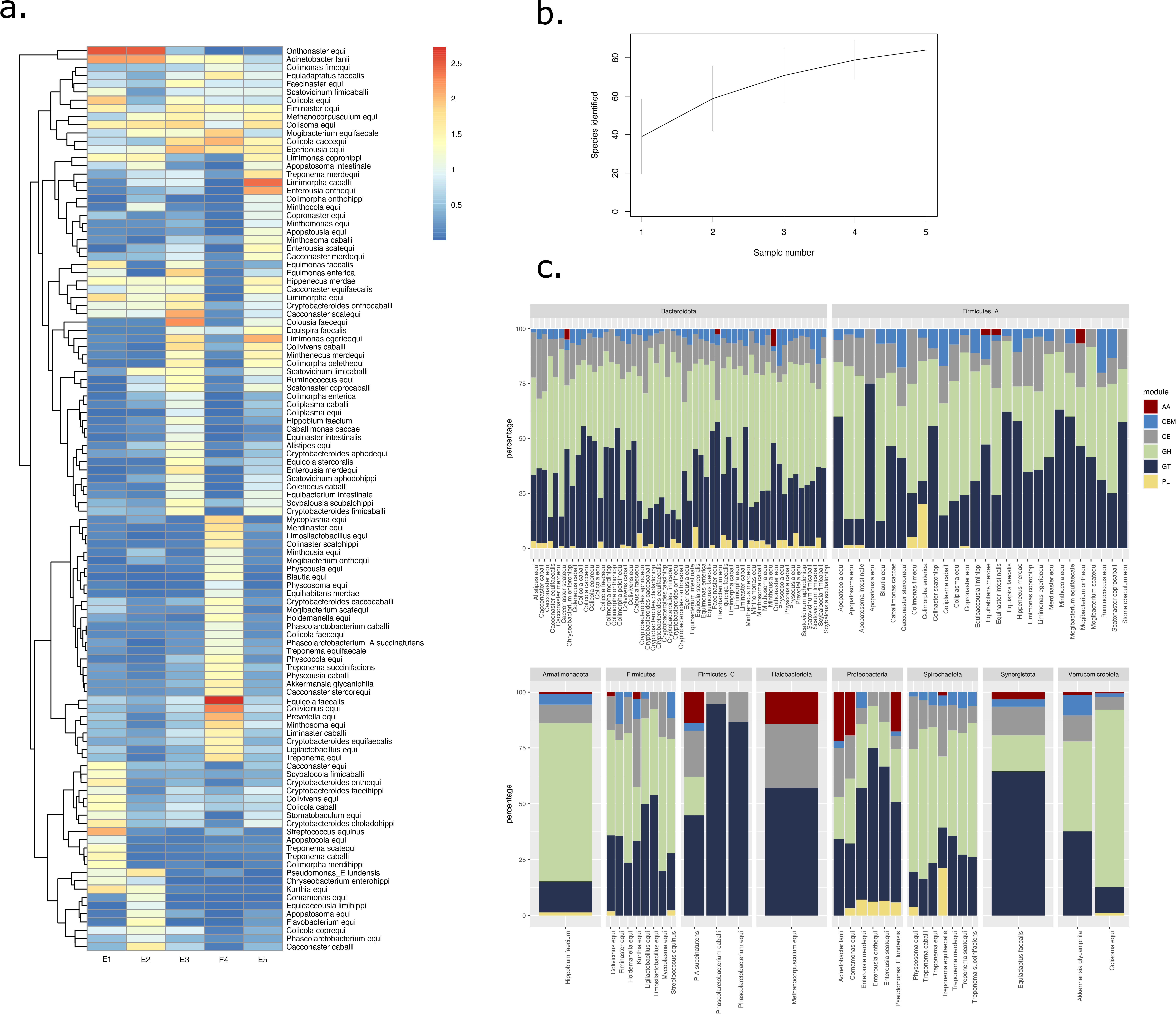
Distribution and metabolism of equine microbial genomes. **(A)**. Heat map depicting the abundance of 110 MAGs across 5 metagenomic samples. Metagenomic species have been annotated with their taxonomic class and species assignment. All data were Log_10_ transformed with Blue colour depicting species of low abundance and Red showing high abundance. **(B)** Species accumulation curve based on coverage data for 110 MAGs over 5 metagenomic samples. (**C)** Proportion of functional CAZyme classes encoded by metagenomic assemblies. Species have been ordered according to GTDB-tk assigned phylum, with functional classes depicted by bar colour.

We created a catalogue of 228,125 genes from our medium- and high-quality MAGs. All 123 MAGs encoded known carbohydrate-active enzymes (CAZymes), with an average of 69 CAZymes per genome (S9 Table). Most (>70%) metagenomic species with a higher-than average repertoire of CAZymes belonged to the *Bacteroidota*. Of the ∼8,500 CAZyme genes reported, most were associated with classes devoted to assembly (glycosyltransferases [GT] 29%) and breakdown (glycoside hydrolases [GH] 51%) of carbohydrate complexes, with far fewer from other groups of CAZymes; being the polysaccharide lyases (PL) and carbohydrate esterases (CE) alongside two further non-enzymatic groups being the carbohydrate-binding modules (CBM) and the auxiliary activities (AA). (Fig 2c). Recovery of 93 classes of glycoside hydrolases from the equine gut mirrors similar enzymes in the sheep rumen linked to fibre degradation (86). Over half of our equine MAGs encode CAZymes with presumed involvement in degradation of hemi-cellulose (58%), cellulose (51%) or pectin or soluble fibre (>60%).

### Many novel bacteriophage genomes

The program VirSorter classified 2,500 contigs as “highly likely” or “likely” to originate from bacteriophages (S10 Table). Of these, 190 bacteriophage genomes were identified as “high-quality” (n=181) or “complete” (n=9) after de-replication (Fig 3a). However, as none showed close identity to known viral sequences, they all represent novel bacteriophage species. Genome sizes ranged from 5 kb to 145 kb, including 42 genomes ranging from 5 to 15 kb in length. Using the viral taxonomy tool Demovir, we could assign 150 of these new phages to known viral families. An additional 29 could be assigned to taxonomically informative viral clusters, based on similarities between predicted proteins from our contigs and proteins from the viral component of the Refseq94 database (S11 Table). Just under half (n=14) of these viral clusters contained at least one reference genome, thus expanding the known diversity of four viral families (Fig 3b).

**Fig 3.**
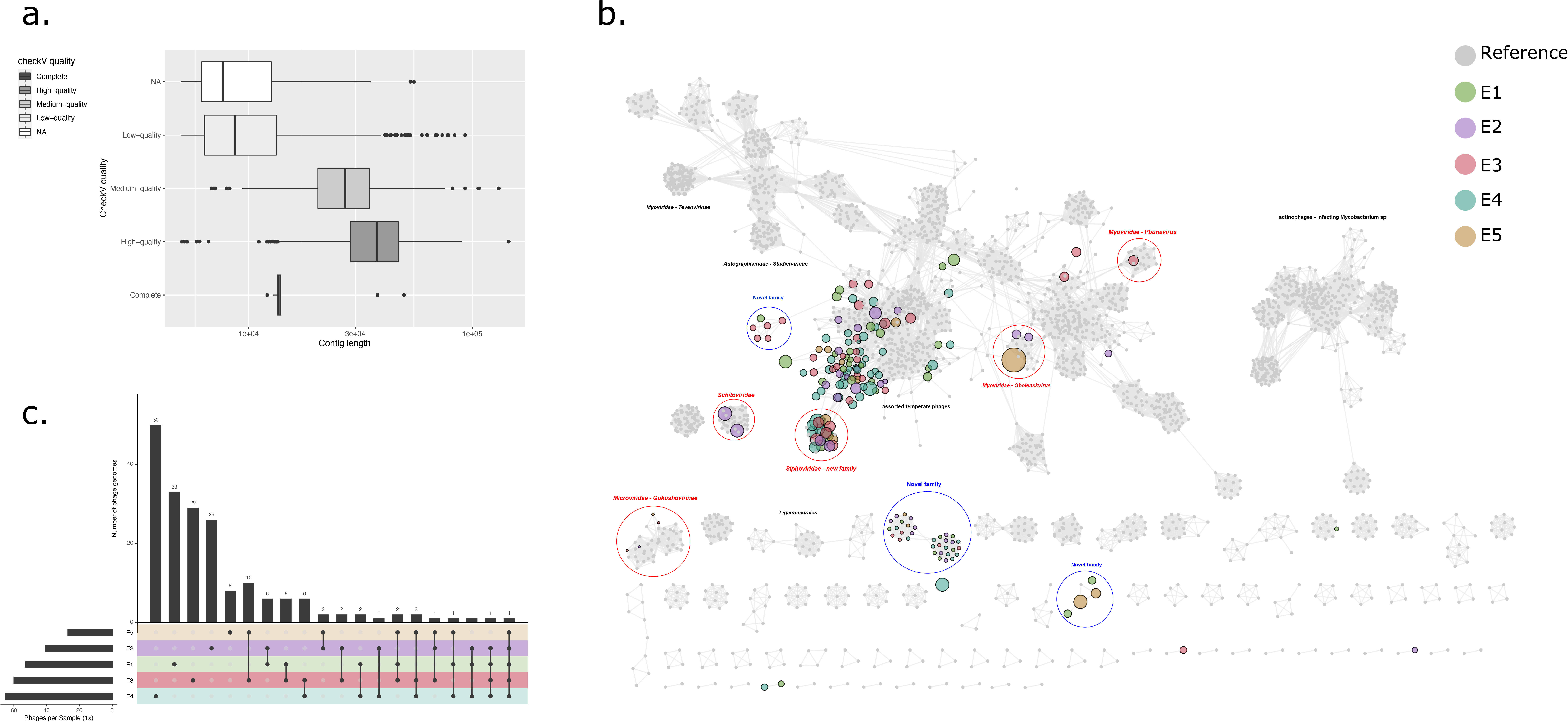
Bacteriophage analysis of equine faecal samples from five thoroughbred horses. (**A)** CheckV quality tiers versus contig length (provided as Log_10_ values). (**B**) Protein sharing network of 190 High-quality or Complete phage genomes assembled from five equine faecal metagenomes and compared against a de-replicated RefSeq database of reference prokaryotic virus genomes. Each node represents a viral genome, with node colour depicting source sample and node size scales according to metagenome contig length. Grey nodes depict reference genomes, with no size scaling shown. Network edges indicate statistically significant relationships between the protein profiles of respective viral genomes. Annotation has been provided to highlight viral clusters of interest. (**C)** Upset plot of phage genomes shared between or specific to source faecal sample, set colour is defined by sample. Each bar represents the number of phage genomes described within the given samples.

Almost all of our viral genomes represented tailed dsDNA phages from the order *Caudovirales* (87) and could be sub-classified into the families *Siphoviridae* (73%), *Podoviridae* or the newly delineated *Schitoviridae* (88) (21%) and *Myoviridae* (6%). Seven genomes were assigned to ssDNA viruses from the family *Microviridae,* four of which cluster as part of the subfamily *Gokushovirinae*. Weak connections of three viral genomes to a viral cluster of *Obolenskvirus,* who’s known members all infect *Acinetobacter* sp., likely indicates the presence of novel bacteriophage genomes predating on the prominent population of bacterial *Acinetobacter* within the equine hind-gut. Present within the viral cluster network but notably absent within our bacteriophage catalogue included the model *Escherichia coli* phages T4 (*Tevenvirinae*) and T7 (*Studiervirinae*) or the *Mycobacterium* infecting actinophages. We observed several novel viral clusters comprising only genomes assembled in this study, which could be classified as the first representatives of new horse hindgut-associated phage families. Based on the proteome comparisons (Fig 3b), we predict at least three new families.

Over three quarters of the recovered phage genomes were found at >1x coverage in just a single sample (Fig 3c). Only one phage was found in all five samples, with coverage ranging from 1.9x - 29x and forming a viral cluster with Lactococcus phage P087 of the family *Siphoviridae*.

## Discussion

Compared to the human gut, the microbiology of the horse gut remains largely unexplored. Here, we deliver new insights into this important ecosystem while also showcasing the advantages of shotgun metagenomics in providing catalogues of genes and genome sequences that take us well beyond what can be achieved using 16S ribosomal RNA gene sequences. Exploration of just five faecal samples allowed discovery of—and recovery of genomes from—over 100 new bacterial and archaeal species and nearly 200 bacteriophage genomes, substantially increasing the known microbial diversity of this environment. Deposition of genomes from these species into publicly available databases will underpin all future studies, improving the quality of reference-based taxonomic assignments.

While the limited scope of this study means it cannot hope to provide a comprehensive view of taxonomic diversity within the horse gut, it gives us a tantalizing glimpse of the richness that awaits us when such approaches are rolled out more widely, particularly as integration of long-read sequencing into metagenomics brings the promise of genome assemblies rivaling those from cultured isolates (89–91). Just as the horse allowed humans to explore new external landscapes, new sequencing and bioinformatics approaches will allow is to explore the inner world of the equine gut microbiome.

## Acknowledgements

The authors would like to thank all horse owners, stud farms and vets for facilitating the collection of horse faecal samples. The authors would also like to thank the core bioinformatics team working at Quadram Institute Bioscience for their help in sample and data processing.

## Author Contributions

RG analysed the data, prepared figures and/or tables, authored or reviewed drafts of the paper, and approved the final draft. JL performed the experiments, authored or reviewed drafts of the paper, and approved the final draft. AR analysed the data, authored or reviewed drafts of the paper, and approved the final draft. EMA analysed the data, authored or reviewed drafts of the paper, and approved the final draft. AO analysed the data, prepared figures and/or tables, authored or reviewed drafts of the paper, and approved the final draft. DB performed the experiments, authored or reviewed drafts of the paper, and approved the final draft. RML & CP conceived and designed the experiments, authored or reviewed drafts of the paper, and approved the final draft. MJP conceived and designed the experiments, analysed the data, prepared figures and/or tables, authored or reviewed drafts of the paper, and approved the final draft

## Data Availability

Data are available on the Sequence Read Archive at BioProject ID PRJNA590977 and Figshare: https://doi.org/10.6084/m9.figshare.14268095

## Supporting information

**Additional File 1. (xlsx.)**

**S1 Table. Sequence Summaries.** Summaries of sequencing data from 5 metagenomic samples sourced from equine faeces from BioProject PRJNA590977

**S2 Table**. **Read-based taxonomic analysis.** Bracken read based relative abundance values for 5 equine faecal samples from BioProject PRJNA590977.

**S3 Table**. **Metagenome Assembled Genome statistics**. CheckM genome statistics for MAG catalogue following assembly of metagenomic reads from 5 horse faecal samples. For High and medium quality sequences, clusters at 95% and 99% ANI have been detailed.

**S4 Table**. **tRNA presence in high and medium quality MAGs**. tRNA presence across MAG catalogue following assembly of metagenomic reads from 5 horse faecal samples. **S5 Table. MAG taxonomic assignments**. Taxonomic analysis for de-replicated metagenomic species according to GTDB (release 95), CAT/BAT (NCBI nr database) and ReferenceSeeker (RefSeq database). Newly assigned Latin binomials have been provided where appropriate.

**S6 Table. AAI analysis for novel genera**. Average amino acid identity (AAI) scores for all genomes of novel genera as determined by CompareM.

**S7 Table. AAI characterisation for novel *Armatimonadota* species**. Reference sequences associated with relevant *Armatimonadota* species according to GTDB and NCBI used for phylogenomic and AAI analysis against our query *Armatimonadota* sequence.

**S8 Table**. **Distribution analysis of recovered MAGs.** Coverage statistics for 110 metagenomic species recovered from 5 metagenomic samples derived from equine faeces. **S9 Table**. **Functional annotation of recovered MAGs**. Presence of genes associated with CAZyme function across 123 MAGs recovered from metagenomic reads of 5 horse faecal samples.

**S10 Table**. **Genome and quality analysis of with recovered phage sequences**. CheckV summary statistics of all VirSorter Category 1 and Category 2 phages >5 kb in length derived from metagenomic assemblies of horse faecal samples. For all High and medium quality phage sequences, further detail of taxonomic annotation and sequences coverage have been provided.

**S11 Table**. **Protein based clustering of high quality or complete phage sequences**. vCONTACT2 output for de-replicated catalogue of 190 phage genomes derived from equine faecal samples and classified as ‘High-quality’ or ‘Complete’ by CheckV

